# ∂Voltage-gated calcium channel activity of gonadotropin-releasing hormone (GnRH) neurons is altered by age and by prenatal androgen exposure in female mice

**DOI:** 10.64898/2026.02.25.707515

**Authors:** Xi Chen, Jennifer Jaime, R. Anthony DeFazio, Suzanne M. Moenter

## Abstract

Polycystic ovary syndrome (PCOS), a common cause of infertility, is marked by persistently high luteinizing hormone (LH)-pulse frequency, presumably driven by high-frequency GnRH pulses. Prenatally androgenized (PNA) mice mimic neuroendocrine PCOS symptoms including high LH-pulse frequency. GnRH neurons from adult PNA mice have a higher firing rate than those from vehicle (VEH) mice; this is reversed in prepubertal mice despite more excitatory inputs at both ages. We hypothesized voltage-gated Ca^2+^ currents (I_Ca_) help set intrinsic excitability of GnRH neurons and are altered by development and/or PNA treatment. Whole-cell patch-clamp recordings were used to measure GnRH neuron I_Ca_ in 3wk-old and adult VEH and PNA mice. PNA treatment increased I_Ca_ density and depolarized the I_Ca_-half inactivation potential at both ages. In VEH but not PNA mice, the Ca^2+^-half activation potential was depolarized in adults versus 3wks. Age decreased the inactivation rate of a fast I_Ca_ regardless of PNA treatment. GnRH neuron firing rate during current injections was higher at 3wks than in adulthood in VEH mice only. Blocking small-conductance Ca²⁺-activated K⁺ current with apamin increased GnRH neuron firing rate except in adult PNA mice. Apamin changed the post-spike-train membrane response from hyperpolarization to depolarization; during development, this net effect of apamin became smaller in PNA mice. In summary, while GnRH neurons from PNA mice have increased I_Ca_, they lack some developmental changes in I_Ca_ kinetics and intrinsic excitability observed in VEH mice. Ca²⁺-activated K⁺ currents are less prominent in GnRH neurons from adult PNA mice, perhaps contributing to increased spontaneous firing.

**Significance statement:** Hyperactivation of GnRH neurons, which control reproductive endocrine function, can lead to increased LH-pulse frequency and is a hallmark of hyperandrogenemia polycystic ovary syndrome (PCOS). We used a mouse model of prenatal androgenization (PNA) that recapitulates the neuroendocrine aspects of PCOS to test the role of calcium currents (I_Ca_) in the PNA phenotype and the typical pubertal process. PNA treatment increased I_Ca_ in GnRH neurons both before and after puberty. Calcium plays a crucial role in neurosecretion thus this may enhance GnRH release. Another role of calcium is activation of calcium-sensitive potassium currents, which tend to decrease action potential firing rate. Despite increased I_Ca,_ calcium-activated potassium currents are less effective in adult PNA mice, perhaps contributing to GnRH neuron hyperactivation.

## Introduction

In vertebrates, pulsatile GnRH secretion from neurons in the preoptic area (POA) and hypothalamus is the ultimate neural mechanism that regulates reproduction. GnRH stimulates secretion of luteinizing hormone (LH) and follicle-stimulating hormone (FSH) from the anterior pituitary (Belchetz et al., 1978; Clarke and Cummins, 1982; Moenter et al., 1992); LH and FSH control steroidogenesis and steroids feed back to regulate GnRH, LH and FSH release. GnRH-pulse frequency varies during the female reproductive cycle and helps drive differential LH and FSH release. GnRH frequency is relatively low in the early follicular phase favoring FSH, and higher in the late follicular phase favoring LH (Wildt et al., 1981; Dalkin et al., 1989; Moenter et al., 1991; McCartney et al., 2022). Disruption of GnRH/LH patterns can lead to infertility as in polycystic ovary syndrome (PCOS), the most common endocrine fertility disorder in females. Hyperandrogenemic PCOS patients exhibit persistently high LH-pulse frequency, presumably driven by high-frequency GnRH pulses, and are resistant to sex steroid negative feedback (Pastor et al., 1998; Eagleson et al., 2000). The high LH-pulse frequency exacerbates ovarian androgen production and can lead to relative FSH deficiency, impairing follicular maturation (Burt Solorzano et al., 2012; McCartney et al., 2022). The hormone dysregulation may begin before or during the pubertal transition, as hyperandrogenemic adolescent girls exhibit higher LH-pulse frequency, and half are resistant to progesterone inhibition (Apter et al., 1994; Chhabra et al., 2005; Blank et al., 2009; Burt Solorzano et al., 2012).

The etiology of PCOS is not fully understood. Besides possible genetic causes, *in utero* androgen exposure could reprogram the fetus and produce PCOS-like characteristics in adulthood (Homburg et al., 2017). Consistent with this postulate, women with PCOS have higher androgen levels during late gestation (Sir-Petermann et al., 2002; Sir-Petermann et al., 2012; Risal et al., 2019) and prenatal androgen (PNA) treatment of several species, including monkeys (Dumesic et al., 1997), sheep (Robinson et al., 2002), rats (Wu et al., 2010) and mice (Sullivan and Moenter, 2004; Moore et al., 2013) results in female offspring with disrupted reproductive cycles, elevated testosterone levels, increased LH-pulse frequency, and resistance to steroid negative feedback. Consistent with elevated LH pulse frequency (Moore et al., 2015), PNA mice exhibit increased spontaneous GnRH neuron action potential firing rate, but interestingly, only in adulthood. In contrast to control female mice, which exhibit a peak of GnRH neuron firing activity at 3wks of age before declining to adult levels, GnRH neuron firing rate is similar throughout development in PNA mice; as a result, firing rate is lower than controls at 3wks of age (Dulka and Moenter, 2017). These changes in firing rate likely represent integration of synaptic and intrinsic changes. In this regard, GABAergic fast synaptic transmission to GnRH neurons, which can be excitatory, is elevated at both 3wks of age and in adults (DeFazio et al., 2002; Sullivan and Moenter, 2004; Moore et al., 2015; Berg et al., 2018). GnRH neuron K^+^ currents properties change with both development and PNA treatment (Jaime et al., 2024), whereas intrinsic excitability of GnRH neurons when intracellular Ca^2+^ is buffered is primarily altered by developmental stage, and not PNA treatment (Jaime and Moenter, 2022).

Voltage-gated Ca^2+^ currents play important roles in mediating excitation-secretion coupling in GnRH neurons (Chen and Moenter, 2023), and regulating GnRH neuron burst firing (Lee et al., 2010). It is unknown if Ca^2+^ currents are altered by development and/or PNA treatment, and if they play a role in shaping GnRH neuron excitability. Here we investigated if there are changes in voltage-gated Ca^2+^ currents in prepubertal vs adult GnRH neurons from control mice, and if PNA treatment alters any Ca^2+^ current properties. We also tested if the small-conductance Ca²⁺-activated K⁺ currents influence GnRH neuron excitability in control and PNA mice before and after puberty.

## Materials and Methods

All chemicals were acquired from Sigma-Aldrich (St. Louis, MO, USA) unless noted otherwise.

### Animals

C57BL/6J mice with green fluorescent protein (GFP) expression targeted to GnRH neurons (GnRH-GFP mice, CBA-Tg(Gnrh1-EGFP)51Sumo/J, MGI:6158457; strain #033639) were bred in our colony (Suter et al., 2000). Mice were housed on a 14h-light 10h-dark photoperiod with lights on at 3am (eastern standard time). Harlan 2916 (non-breeders) or 2919 (breeders) chow and water were available *ad libitum*. To generate PNA mice, a homozygous GnRH-GFP female was bred with a C57BL/6J male (Jackson Labs 000664). A companion CD1 female was also in the cage for breeding to provide maternal care and nutrition. The first day of pregnancy in female was determined by the appearance of a copulatory plug during daily checks. On day 16 to day 18 of pregnancy, the GnRH-GFP dam received 225μg dihydrotestosterone (DHT) in 50µl sesame oil sc for PNA treatment, or sesame oil vehicle (VEH) for controls. Mice were weaned at PND 21. CD1 pups, identified by coat color, were culled as needed to avoid crowding. All procedures were approved by the Institutional Animal Care and Use Committee of the University of Michigan.

### Verification of PNA phenotype

The PNA status was confirmed by the age of vaginal opening (VO), the anogenital distance (AGD) measured at PND 70-72, and estrous cyclicity via vaginal lavage monitored at PND 70-83. For mice recorded before puberty, the PNA status was confirmed by phenotypes of littermates raised to adulthood, as these phenotypes are consistent among littermates.

### Brain slice preparation

Brain slices were prepared as described (Jaime et al., 2024). All bath solutions were bubbled with 95% O_2_/5% CO_2_ for at least 30min before exposure to tissue and throughout experiments. The mouse brain was quickly removed and placed in the ice-cold sucrose saline containing 250mM sucrose (Invitrogen), 3.5mM KCl (Fluka), 26mM NaHCO_3_, 10mM D-glucose, 1.25mM Na_2_HPO_4_⋅7H_2_O (Mallinkrodt), 1.2mM MgSO_4_⋅7H_2_O (EMD Millipore), and 2.5mM MgCl_2_ (Fluka), at pH 7.6 and 345mOSM. Coronal brain slices (300μm) through the POA of hypothalamus were cut using a VT1200 S vibratome (Leica Biosystems). Brain slices were kept in 1:1 mixture of sucrose saline and artificial cerebrospinal fluid (ACSF) containing 135mM NaCl (Fluka), 3.5mM KCl, 26mM NaHCO_3_, 1.25mM Na_2_HPO_4_⋅7H_2_O, 2.5mM CaCl_2_, 1.2mM MgSO_4_⋅7H_2_O, and 10mM D-glucose at pH 7.4 and 305mOSM, at room temperature for 30min. Brain slices were then transferred to 100% ACSF and incubated for 0.5-6h before recording. No differences were noted in data based upon time after slice preparation within this range.

### Recording solutions and data acquisition

During recording, the brain slice was placed in a chamber and consistently perfused (1.5-2ml/min) with solutions (see below) kept between 29–31°C with an inline heater (Warner Instruments, Hamden, CT). GFP-positive GnRH neurons were identified by a brief illumination at 470nm using an Olympus BX51WI microscope. Whole-cell voltage-clamp and current-clamp recordings were made using one channel of a HEKA dual EPC10 amplifier running PatchMaster (HEKA Elektronik). Data were sampled at 10kHz and filtered at 7.3kHz.

For recording Ca^2+^ currents, brain slices were perfused with a solution containing: 120mM NaCl, 10mM glucose, 26mM NaHCO_3_, 1.25mM Na_2_HPO_4_, 1.2mM MgSO_4_, 1.0mM CaCl_2_, 1.5mM MgCl_2_, 5mM 4-aminopyridine (4-AP), 5mM CsCl, 10mM TEA-Cl, 2µM TTX, pH 7.7 and 305 mOSM. Pulled capillary glass pipette (3-3.9MΩ) were filled with a solution containing 120mM Cs-gluconate, 10mM HEPES, 10mM EGTA, 0.5mM CaCl_2_, 4mM Mg-ATP, 0.4mM Na-GTP, 20mM TEA-Cl, pH 7.2 and 301 mOSM. The pipette was placed at the surface of a GnRH neuron soma and a seal of >1GΩ was formed with the cell membrane. After stabilizing for 3min, a gentle suction was applied to achieve the whole-cell configuration. Cells were stabilized in the whole-cell configuration for 30s to allow dialysis of the pipette solution and were held at -70mV between protocols. Slow capacitive current was electronically compensated, and R_s_ was auto-compensated with prediction of 50%-75%. Recordings were corrected online for a liquid junctional potential of -13.3mV. Leak current was subtracted online (P/-10) using an average of 6 sweeps from a baseline potential of -70mV (Bezanilla and Armstrong, 1977). Before and after a series of voltage steps, input resistance (R_in_), series resistance (R_s_), holding current (I_hold_) and capacitance were measured in voltage-clamp mode from the mean current response to twenty 5mV-hyperpolarizing steps from -70mV. Cells had high R_in_ attributable to the blockage of potassium channels. Recordings were analyzed only if the following quality parameters were met: Cm 8 to 20pF, R_in_ >900MΩ, stable R_s_ <22MΩ, I_hold_ of -55 to 0pA. Pilot studies were conducted to examine rundown of peak current and based on these all recordings were completed within four min of achieving the whole-cell configuration.

Action potentials during current injections were recorded to characterize excitability. To evaluate the excitability of GnRH neurons without extrinsic calcium buffering, pipettes were filled with EGTA-free solution containing 136.5mM K gluconate, 20.4mM NaCl, 10.5mM HEPES, 4.2mM MgATP, 0.4mM NaGTP, 10.3mM KOH, pH 7.2 with NaOH, 310mOSM. Brain slices were perfused with ACSF containing 20µM D-APV (Tocris), 10µM CNQX (Tocris) and 100µM picrotoxin to block ionotropic glutamate and GABA receptors. After achieving the whole-cell configuration as described above, series resistance was calculated during slow-capacitance compensation for an estimation of bridge balance, and the recording mode was transferred to current-clamp with a bridge balance of 95%. Additional direct current (<5pA) was injected if needed to keep the baseline membrane potential near -70.7±0.1mV (range: -72.9 to -68.4mV). Recordings were corrected online for a liquid junctional potential of -14.4mV and were analyzed only if the following quality parameters were met: Cm 8 to 20pF, R_in_ >500MΩ, stable R_s_ <22MΩ, I_hold_ of -30 to 10pA.

### Experimental design

Calcium currents and excitability of GnRH neurons were measured in prepubertal (PND 18-21, before weaning) and adult mice (PND 84-155). Mice were treated prenatally with either VEH or DHT resulting in four groups: 3wk VEH, 3wk PNA, adult VEH and adult PNA. For each group, a minimum of five mice from at least four litters were used. No more than three GnRH neurons from the same mouse were analyzed.

### Voltage-gated Ca^2+^ current characterization

Ca^2+^ currents were isolated pharmacologically by using nominally K^+^-free pipette and external solutions and blocking voltage-gated K^+^ (4-AP and TEA-Cl) and Na^+^ (TTX) channels. Pilot studies indicated that depolarization at -40mV for 200ms inactivates the fast-inactivating component of the Ca^2+^ current, and hyperpolarization at -100mV for 2000ms completely removes inactivation. Fast and medium/slow-inactivating components of Ca^2+^ current (fast I_Ca_ and medium/slow I_Ca_, respectively) were thus distinguished based on voltage dependence and time course.

### Voltage-dependent activation of Ca^2+^ currents

To measure total Ca^2+^ current, the membrane potential was hyperpolarized to -100mV for 2000ms to remove inactivation, followed by an additional 200ms prepulse at -100mV. The cell was then given a series of 200ms voltage steps from -100mV to +60mV in 10mV increments. After this series, the same cell received the same voltage step protocol, but with the 200ms prepulse at -40mV (rather than -100mV) to inactivate the fast I_Ca_ to reveal the medium/slow I_Ca_ components. The fast I_Ca_ was mathematically isolated by subtracting the medium/slow I_Ca_ obtained with the -40mV prepulse from total current obtained with the -100mV prepulse.

### Voltage-dependent inactivation of fast I_ca_

Depolarization to 0mV produced the largest macroscopic Ca^2+^ current in GnRH neurons and was thus used as the test pulse to evaluate voltage-dependent inactivation of fast I_Ca_. After hyperpolarization to -100mV to remove inactivation, the GnRH neuron was subjected to a 200ms prepulse from -90mV to -40mV at 5mV or 10 mV increments, followed by a test pulse at 0mV for 25ms; data from the two increments were consistent and combined. The more depolarized prepulses lead to smaller current triggered by the 0mV test pulse, reflecting voltage-dependent inactivation of fast I_Ca_. The fast I_Ca_ was isolated by subtracting the test current at 0mV following the -40mV prepulse from test currents with prepulses of -90 to -45mV.

### Time-dependent inactivation and recovery of fast I_ca_

To measure the time course of inactivation of the fast I_Ca_, GnRH neurons were hyperpolarized to -100mV for 2000ms to remove inactivation and then depolarized to the inactivation potential of -40mV for 1, 2, 4, 8, 16, 32, 64, 128, and 256ms, followed by a test pulse at 0mV for 25ms. The non-inactivating component during the test pulse after the 256ms inactivation was subtracted from currents of other sweeps with shorter inactivation times to isolate the fast I_Ca_. To measure the time course of recovery from inactivation of fast I_Ca_, GnRH neurons were first depolarized to -40mV for 200ms to inactive this component. The cell was then subjected to -100mV hyperpolarization for 0, 1, 2, 4, 8, 32, 64, 128, 256, 512, 1024, and 2048ms to recover the fast I_Ca_, followed by a test pulse at 0mV for 25ms. The current during the test pulse without a recovery prepulse (0ms) was subtracted from currents of other sweeps with longer time of recovery to isolate the fast I_Ca_.

### Intrinsic excitability of GnRH neuron

To evaluate GnRH neuron intrinsic excitability, the firing response to sequential current injections (0-100pA, 10pA increment, 500ms) was recorded using whole-cell current-clamp without EGTA buffering of the pipette solution so that the potential influence of Ca^2+^ influx and/or release from intracellular stores was not hindered; additional cells were recorded with EGTA buffering for pilot comparisons. Under EGTA-free conditions, GnRH neurons presented a prolonged, slow hyperpolarization following the termination of current injection, which gradually returned to baseline potential over several seconds. The interval between current steps was thus set to 15s so the membrane potential would return to approximately -70mV before the next step was initiated.

### The effect of small conductance Ca²⁺-activated K⁺ current on GnRH neuron excitability

Apamin (300nM, Tocris) was applied to block the small-conductance Ca^2+^-activated K^+^ channels to investigate the role of these channels in GnRH neuron excitability. Action potentials were first recorded under basal state (0min) in response a train of 500ms-current injections (0, 20, 40, 80pA) delivered at 15s intervals. Then, the brain slice was perfused with apamin or vehicle control and the response to identical current injections were recorded 2, 4, and 6min after the start of apamin or VEH perfusion. Passive properties of the GnRH neuron were recorded in response to the average of twenty 5mV hyperpolarizing steps from -70mV before and after each train of current injections to verify recording stability. UCL 2077(10μM), a blocker of KCNQ1 and KCNQ2 potassium channels (Soh and Tzingounis, 2010), was tested in pilot studies but had no discernable effect on action potential firing under these conditions (data not shown).

### Analysis

For Ca^2+^ current analysis, the current at each voltage step was normalized by cell capacitance to yield current density. To make I-V curves, current densities of total I_Ca_, medium/slow I_Ca_, and fast I_Ca_ were plotted against each command voltage. Because of the potential influence of unblocked potassium currents from 30mV to 60mV, statistical analysis for current density and voltage-dependent activation were limited to the range of -100 to 20mV.

To assess the voltage-dependent activation of total, fast, and medium/slow Ca^2+^ current, Ca^2+^ conductance was calculated by dividing the current peak of each voltage step by the driving force, which is derived from Goldman-Hodgkin-Katz (GHK) equation:

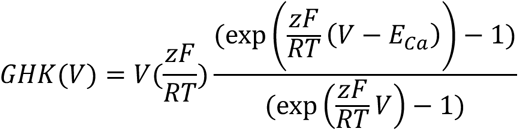

Where V is the step potential, R is the universal gas constant, T is the temperature in Kelvin, z is the valence of Ca^2+^, F is the Faraday’s constant, E_Ca_ was calculated to be 151.43mV.

The Ca^2+^ conductance was normalized to the maximum value and plotted as a function of potential to reveal voltage-dependence of activation. The activation curve was fitted with the Boltzmann equation:

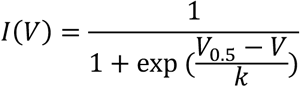

V_0.5_ is the potential that the conductance reaches the half maximum (V_0.5 act_), and k is the slope factor. V_0.5 act_ and k were used to assess the activation of Ca^2+^ current.

To assess the voltage-dependent inactivation of fast I_Ca_, the peak of fast I_Ca_ obtained at 0mV with varying prepulses was normalized to maximum fast I_Ca_, defined as the average of currents following -80, -75 and -70mV prepulses, as voltage-dependent inactivation did not occur in this range. The normalized fast I_Ca_ was plotted against the corresponding command voltage step and fitted to Boltzmann equation as above; V_0.5 inact_ and k were used to assess the inactivation of fast I_Ca_.

To assess the time-dependent inactivation of fast I_Ca_, the peak fast I_Ca_ after 1 to 256ms inactivation prepulses was normalized to the maximum value after 1ms of inactivation, and plotted as a function of inactivation time. The time-dependent inactivation curve was fitted to the monophasic decay equation:

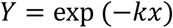

Where k is the rate constant. The time at which half of the fast I_Ca_ was inactivated by -40mV depolarization (T_0.5 inact_) is computed as ln(2)/k.

To assess the time-dependent recovery of fast I_Ca_ after inactivation, the peak fast I_Ca_ after 1 to 2048ms recovery prepulses was normalized to the maximum value after 2048ms of recovery, and plotted as function of recovery time. The time-dependent recovery curve was fitted to the monophasic association equation:

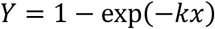

Where k is the rate constant and T_0.5 recov_ was computed as ln(2)/k.

To assess GnRH neuron excitability, firing rate was plotted as a function of current injection amplitude. The first spike fired at rheobase was used to profile action potential characteristics. Action potential latency from the start of current injection, threshold (membrane potential at which the second derivative is >5000mV/s/s), rate of rise, peak amplitude relative to threshold, full width at half-maximum (FWHM), time and amplitude of the afterhyperpolarization potential (AHP) relative to threshold were compared among groups.

Under basal conditions, all GnRH neurons exhibited hyperpolarization after a train of action potentials triggered by the current injection. To analyze the fast and slow post-spike-train membrane potential change relative to the pre-stimulus baseline potential (ΔV_m_), the point at which the repolarization rate after current injection termination fell below 50mV/s was used; the period before being designated fast ΔV_m_ and that after slow ΔV_m_. Peak values of fast and slow ΔV_m_ were compared among groups. The area under the curve (AUC) of ΔV_m_ during the post-stimulus 12.5s period were compared.

To compare the post-spike-train membrane depolarization that emerged with 6min apamin treatment, the post-spike-train depolarization peak and the latency to the peak following termination of current injection were compared. AUC and duration of the post-spike-train depolarization were quantified. To minimize the influence of noise and prolonged tails, the onset and termination were defined as the points when potential crossed 10% of the depolarization peak value (AUC at 10% maximum, duration at 10% maximum).

To assess the net effect of apamin-sensitive SK current on post-spike-train membrane potential, the membrane potential trace recorded starting 6min into apamin treatment was subtracted from that recorded before treatment in the same cell. The SK-mediated hyperpolarization peak value, peak latency, AUC at 10% maximum and duration at 10% maximum were analyzed.

### Statistics

Statistics were performed by Igor 9 (Sutter) and Prism 9 (GraphPad LLC.). Data were tested for normality using the Shapiro-Wilk test. Equality of variance was tested by F test. Data were reported as the mean±SEM (normal distribution) or median±IQR (non-normal distribution). Details of numbers of cells and animals, and of statistical tests are provided in the results and tables

## Results

### Verification of prenatal androgenization phenotype

PNA-induced differences were confirmed by the age of VO, AGD, and estrous cyclicity. Consistent with previous observations (Sullivan and Moenter, 2004; Dulka and Moenter, 2017; Jaime and Moenter, 2022; Jaime et al., 2024), PNA mice exhibited earlier VO (Figure 1A, two-tailed Mann-Whitney U test, U=265.6, p<0.0001), which occurred at a lower body mass (Figure 1B, two-tailed Mann-Whitney U test, U=157, p<0.0001). At PND70-72, PNA mice had increased anogenital distance (AGD, Figure 1C, two-tailed unpaired Welch’s t-test, t=19, df=98, p<0.0001), while body mass did not differ from VEH (two-tailed Mann-Whitney U test, U 1171). Adult PNA mice had disrupted estrous cycles, with more days in diestrus and fewer days in proestrus and estrus compared with controls (Figures 1E, F, Chi-square test p<0.0001, Ξ^2^=190.8, df=2).

**Figure 1.**
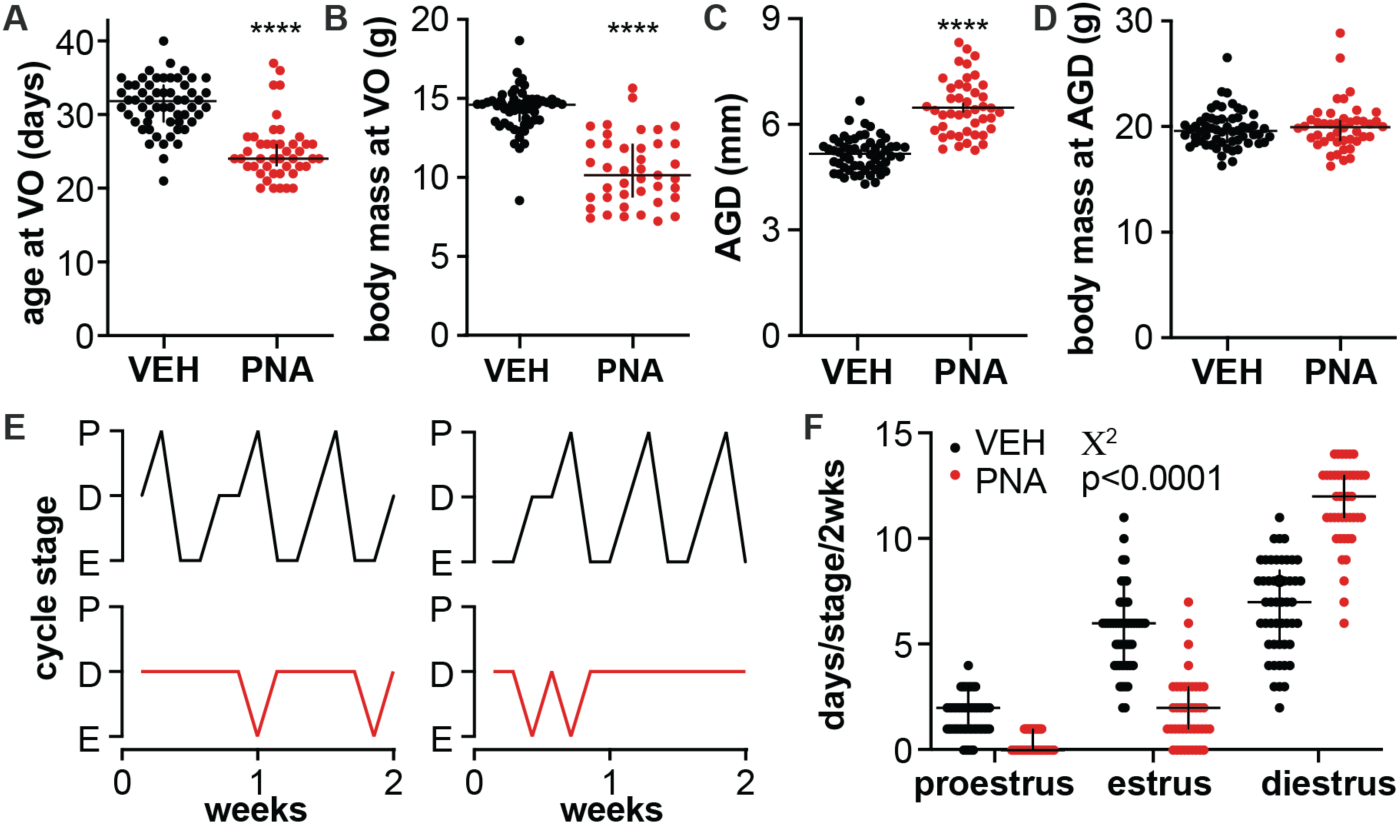
Verification of PNA phenotype. **A, B, D**, Individual values and median±interquartile range for age at vaginal opening (VO) (A), body mass at the day of VO (B), and body mass at PND 70-72 (D). **C**, Individual values and mean±SEM anogenital distance (AGD) measured at postnatal day (PND) 70-72. **E**, Representative estrous cycles of two VEH (top, black) and two PNA (bottom, red) mice over 14 days. P, proestrus; D, diestrus; E, estrus. (F), Days in proestrus, estrus and diestrus stages over 14 days. ****p<0.0001, two-tailed Mann-Whitney U test (A, B, D), two-tailed unpaired Welch’s t-test (C), Ξ^2^ (F).

### Ca^2+^ current density in GnRH neurons is altered by PNA treatment

Ca^2+^ currents were recorded using whole-cell voltage-clamp. Passive properties and series resistance of voltage-clamp recordings were used to assess the recording quality and cell properties. Because potassium channels were blocked, all cells exhibited high input resistance (R_in_, >900MΩ). The cell capacitance was higher in the adult groups compared to 3wk-old groups, but PNA treatment had no effect (two-way ANOVA age p<0.0001, Table 1). There were no differences in R_in_, R_s_, or I_hold_ among groups.

**Table 1.** descriptive statistics and two-way ANOVA parameters for passive properties and recording quality for Ca^2+^ currents recording (Figure 2, 3, 4). **Bold** indicates p<0.05.

| Descriptive statistics |  |  |  |  |
| --- | --- | --- | --- | --- |
| property | 3wk VEH | 3wk PNA | adult VEH | adult PNA |
| series resistance (MΩ) | 14.869 ± 0.481 | 14.642 ± 0.465 | 15.810 ± 0.466 | 15.402 ± 0.490 |
| input resistance (MΩ) | 1634.412 ± 78.424 | 1545.381 ± 73.494 | 1684.143 ± 84.791 | 1530.916 ± 61.880 |
| capacitance (pF) | 12.943 ± 0.444 | 13.882 ± 0.405 | 16.258 ± 0.480 | 16.181 ± 0.484 |
| holding current (pA) | -31.598 ± 1.471 | -32.065 ± 1.537 | -28.319 ± 1.381 | -30.591 ± 1.913 |
| Two-way ANOVA |  |  |  |  |
|  | age | treatment | interaction |  |
| series resistance (MΩ) | F(1,119)= 3.193<br>p=0.0765 | F(1,119)=0.4455<br>p=0.5058 | F(1,119)= 0.03638<br>p=0.8491 |  |
| input resistance (MΩ) | F(1,119)=0.05504<br>p=0.8149 | F(1,119)= 2.597<br>p=0.1097 | F(1,119)= 0.1824<br>p=0.6701 |  |
| capacitance (pF) | **F(1,119)= 38.03**<br>**p<0.0001** | F(1,119)= 0.8962<br>p=0.3457 | F(1,119)= 1.224<br>p=0.2669 |  |
| holding current (pA) | F(1,119)= 2.229<br>p=0.1381 | F(1,119)= 0.7461<br>p=0.3894 | F(1,119)= 0.5700<br>p=0.3244 |  |
| Post-hoc Fisher's LSD analysis of significant two-way ANOVA |  |  |  |  |
|  | 3wk VEH<br>vs.<br>3wk PNA | adult VEH<br>vs.<br>adult PNA | 3wk VEH<br>vs.<br>adult VEH | 3wk PNA<br>vs.<br>adult PNA |
| capacitance (pF) | p=0.1491 | p=0.9047 | **p<0.0001** | **p<0.0001** |

The total I_Ca_ in GnRH neurons is comprised of two main components: a fast-inactivating component (fast I_Ca_) that was inactivated by a 200ms prepulse at -40mV, and a medium/slow inactivating component (medium/slow I_Ca_) that remains after the -40mV inactivation. Representative traces of total I_Ca_, medium/slow I_Ca_ and mathematically-isolated fast I_Ca_ are shown at the top of Figure 2A-C, with current density-voltage (I-V) curves in the middle and current densities at 0mV below. There was an overall effect of PNA treatment on the I-V curves of the total I_Ca_ and each individual component (three-way ANOVA treatment p<0.05, treatment x voltage p<0.0001, Table 2). At 0mV, where the maximum total I_Ca_ was observed, GnRH neurons from PNA mice exhibited higher total I_Ca_ density compared to VEH groups at both 3wks of age and in adulthood. (Figure 2A, bottom, two-way ANOVA treatment p=0.0039, Fisher’s LSD, PNA vs VEH p<0.05 in both age groups, Table 3). This change in total I_Ca_ was attributable to higher densities of both medium/slow and fast I_Ca_ in cells from PNA mice (Figure 2B, C, two-way ANOVA treatment p<0.05 in medium/slow I_Ca_ and fast I_Ca,_ Table 3). Although three-way ANOVA showed the I-V curve of the fast component was affected by age (age x voltage p=0.0259), the *post hoc* analysis revealed the main differences were from +30 to +60mV. At this voltage range, there is concern about the influence of unblocked potassium currents. This statistical difference is thus likely not biologically relevant and is not considered further.

**Figure 2.**
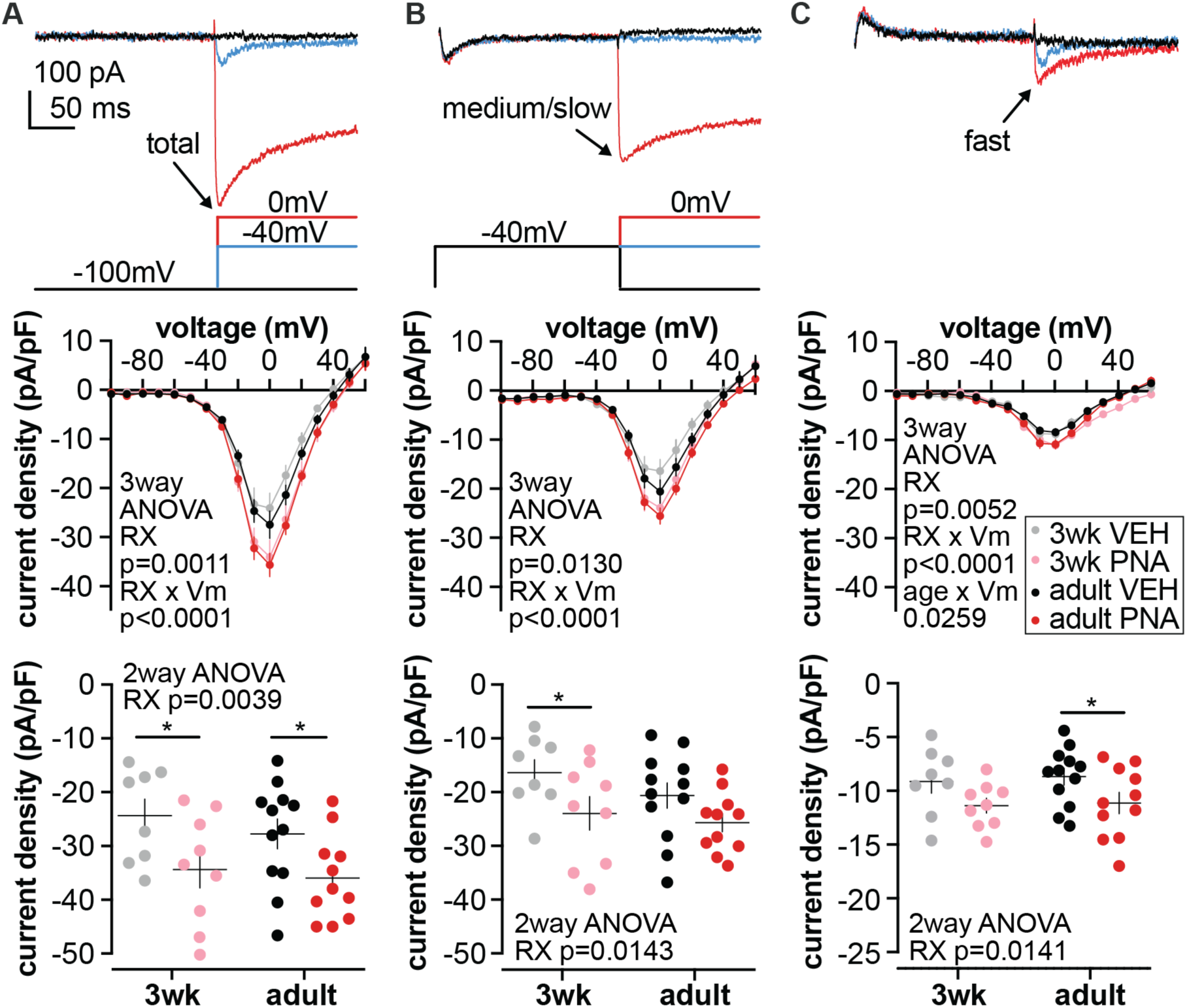
Ca^2+^ currents of GnRH neurons in VEH and PNA mice during development. **A-C** Top: Representative current traces (above) and voltage commands (below) for total I_Ca_ (A), medium/slow I_Ca_ (B), and mathematically-isolated fast I_Ca_ (C) from an adult PNA mouse. Labeled arrows (total, med/slow, fast) identify current components. The 2000ms prepulse at -100mV to remove inactivation is not shown. Only three voltage steps (-100, -40, 0mV) from the voltage family from -100 to 60mV are shown for clarity. Middle: Current density-voltage (I-V) curve for total I_Ca_ (A), medium/slow I_Ca_ (B), and mathematically-isolated fast I_Ca_ (C). Note the Y axis does not cross the X axis at zero. Bottom: current densities of total I_Ca_ (A), medium/slow I_Ca_ (B), and mathematically-isolated fast I_Ca_ (C) at 0mV. *p<0.05, Fisher’s LSD. RX, prenatal treatment.

**Table 2.** Three-way ANOVA and Fisher’s LSD test of current density-voltage (I-V) curve for total I_Ca_, medium/slow I_Ca_, and mathematically-isolated fast I_Ca_ (Figure 2). **Bold** indicates p<0.05.

| source of variation | total $I_{\text{Ca}}$ | medium/slow $I_{\text{Ca}}$ | fast $I_{\text{Ca}}$ |
| --- | --- | --- | --- |
| voltage | **$F(16, 576) = 317.6$**<br>**$p < 0.0001$** | **$F(16, 576) = 215.4$**<br>**$p < 0.0001$** | **$F(16, 576) = 196.5$**<br>**$p < 0.0001$** |
| age | $F(1, 36) = 0.6734$<br>$p = 0.4173$ | $F(1, 36) = 1.450$<br>$p = 0.2363$ | $F(1, 36) = 1.217$<br>$p = 0.2772$ |
| treatment | **$F(1, 36) = 12.50$**<br>**$p = 0.0011$** | **$F(1, 36) = 6.838$**<br>**$p = 0.0130$** | **$F(1, 36) = 8.875$**<br>**$p = 0.0052$** |
| voltage x age | $F(16, 576) = 0.6167$<br>$p = 0.8717$ | $F(16, 576) = 1.219$<br>$p = 0.2475$ | **$F(16, 576) = 1.818$**<br>**$p = 0.0259$** |
| voltage x treatment | **$F(16, 576) = 6.883$**<br>**$p < 0.0001$** | **$F(16, 576) = 4.335$**<br>**$p < 0.0001$** | **$F(16, 576) = 4.010$**<br>**$p < 0.0001$** |
| age x treatment | $F(1, 36) = 0.08796$<br>$p = 0.7685$ | $F(1, 36) = 0.04491$<br>$p = 0.8334$ | $F(1, 36) = 0.1523$<br>$p = 0.6986$ |
| voltage x age x treatment | $F(16, 576) = 0.2674$<br>$p = 0.9982$ | $F(16, 576) = 0.4529$<br>$p = 0.9673$ | $F(16, 576) = 1.756$<br>$p = 0.0338$ |

| Table 2 continued |  |  |  |  |
| --- | --- | --- | --- | --- |
| Post-hoc Fisher's LSD analysis of three-way ANOVA for total I<sub>Ca</sub> current densities at different voltage steps |  |  |  |  |
| test potential (mV) | 3wk VEH<br>vs.<br>3wk PNA | adult VEH<br>vs.<br>adult PNA | 3wk VEH<br>vs.<br>adult VEH | 3wk PNA<br>vs.<br>adult PNA |
| -100 | p=0.9481 | p=0.8626 | p=0.9593 | p=0.9693 |
| -90 | p=0.9417 | p=0.8371 | p=0.8979 | p=0.6890 |
| -80 | p=0.8411 | p=0.9061 | p=0.9864 | p=0.9284 |
| -70 | p=0.8163 | p=0.9898 | p=0.9310 | p=0.8610 |
| -60 | p=0.8733 | p=0.9506 | p=0.9731 | p=0.7917 |
| -50 | p=0.8190 | p=0.9795 | p=0.9563 | p=0.8293 |
| -40 | p=0.8854 | p=0.8517 | p=0.9959 | p=0.7379 |
| -30 | p=0.7541 | p=0.4474 | p=0.8103 | p=0.9019 |
| -20 | p=0.0684 | **p=0.0094** | p=0.4439 | p=0.7410 |
| -10 | **p=0.0003** | **p<0.0001** | p=0.4813 | p=0.4955 |
| 0 | **p<0.0001** | **p<0.0001** | p=0.0882 | p=0.4195 |
| 10 | **p<0.0001** | **p=0.0006** | **p=0.0472** | p=0.5132 |
| 20 | **p=0.0011** | **p=0.0120** | p=0.1471 | p=0.7948 |

| Post-hoc Fisher's LSD analysis of three-way ANOVA for medium/slow I<sub>Ca</sub> current densities at different voltage steps |  |  |  |  |
| --- | --- | --- | --- | --- |
| test potential (mV) | 3wk VEH<br>vs.<br>3wk PNA | adult VEH<br>vs.<br>adult PNA | 3wk VEH<br>vs.<br>adult VEH | 3wk PNA<br>vs.<br>adult PNA |
| -100 | p=0.9771 | p=0.8655 | p=0.9000 | p=0.9515 |
| -90 | p=0.9755 | p=0.7733 | p=0.9507 | p=0.7162 |
| -80 | p=0.8678 | p=0.7857 | p=0.9237 | p=0.7374 |
| -70 | p=0.9768 | p=0.6780 | p=0.9262 | p=0.7948 |
| -60 | p=0.9973 | p=0.7459 | p=0.9008 | p=0.8645 |
| -50 | p=0.9827 | p=0.9250 | p=0.9681 | p=0.9167 |
| -40 | p=0.6532 | p=0.7691 | p=0.5451 | p=0.8857 |
| -30 | p=0.9530 | p=0.5432 | p=0.5095 | p=0.9670 |
| -20 | p=0.1305 | **p=0.0250** | p=0.6322 | p=0.9714 |
| -10 | **p=0.0009** | **p=0.0022** | p=0.2257 | p=0.6355 |
| 0 | **<0.0001** | **p=0.0016** | **p=0.0151** | p=0.3169 |
| 10 | **p=0.0012** | **p=0.0058** | **p=0.0465** | p=0.2809 |
| 20 | **p=0.0169** | p=0.0864 | p=0.0761 | p=0.4180 |

| Post-hoc Fisher's LSD analysis of three-way ANOVA for fast I<sub>Ca</sub> current densities at different voltage steps |  |  |  |  |
| --- | --- | --- | --- | --- |
| test potential (mV) | 3wk VEH<br>vs.<br>3wk PNA | adult VEH<br>vs.<br>adult PNA | 3wk VEH<br>vs.<br>adult VEH | 3wk PNA<br>vs.<br>adult PNA |
| -100 | p=0.5560 | p=0.7287 | p=0.5560 | p=0.3613 |
| -90 | p=0.9900 | p=0.5853 | p=0.9255 | p=0.6904 |
| -80 | p=0.5536 | p=0.8129 | p=0.6744 | p=0.9953 |
| -70 | p=0.3782 | p=0.8463 | p=0.4241 | p=0.7482 |
| -60 | p=0.1840 | p=0.9244 | p=0.5325 | p=0.3728 |
| -50 | p=0.2967 | p=0.3317 | p=0.5524 | p=0.1539 |
| -40 | p=0.9623 | p=0.8076 | p=0.6234 | p=0.5006 |
| -30 | p=0.5374 | p=0.3925 | p=0.5811 | p=0.4916 |
| -20 | **p=0.0248** | p=0.0337 | p=0.6534 | p=0.3616 |
| -10 | p=0.1148 | **p=0.0002** | p=0.4696 | p=0.3164 |
| 0 | **p=0.0056** | **p=0.0004** | p=0.5633 | p=0.7524 |
| 10 | **p=0.0011** | **p=0.0229** | p=0.4081 | p=0.5595 |
| 20 | **p=0.0074** | p=0.0509 | p=0.7012 | p=0.1403 |

**Table 3.** Two-way ANOVA of current density at 0mV (Figure 2).

| current type | age | treatment |  | interaction |
| --- | --- | --- | --- | --- |
| total I<sub>Ca</sub> | F (1, 36) = 0.7165<br>p=0.4029 | F (1, 36) = 9.540<br>p=0.0039 |  | F (1, 36) = 0.09465<br>p=0.7601 |
| medium/slow I<sub>Ca</sub> | F (1, 36) = 1.475<br>p=0.2324 | F (1, 36) = 6.625<br>p=0.0143 |  | F (1, 36) = 0.2647<br>p=0.6100 |
| fast I<sub>Ca</sub> | F (1, 36) = 0.1373<br>p=0.7131 | F (1, 36) = 6.653<br>p=0.0141 |  | F (1, 36) = 0.01243<br>p=0.9119 |
| Post-hoc Fisher's LSD analysis of significant two-way ANOVA |  |  |  |  |
|  | 3wk VEH<br>vs.<br>3wk PNA | adult VEH<br>vs.<br>adult PNA | 3wk VEH<br>vs.<br>adult VEH | 3wk PNA<br>vs.<br>adult PNA |
| total I<sub>Ca</sub> | p=0.0314 | p=0.0397 | p=0.4233 | p=0.7033 |
| medium/slow I<sub>Ca</sub> | p=0.0492 | p=0.1228 | p=0.2329 | p=0.6208 |
| fast I<sub>Ca</sub> | p=0.1125 | p=0.0462 | p=0.7371 | p=0.8545 |

### Voltage-dependent activation of the Ca^2+^ conductance in GnRH neurons is affected by development in VEH mice

Ca^2+^ conductance was calculated by dividing the peak current at each voltage step by the driving force derived from GHK equation to assess voltage-dependent activation. The Ca^2+^ conductance at each voltage step was normalized to the maximum current value, typically achieved at 10 or 20mV, and were plotted as a function of test potential (Figure 3A, B, C, top). There was an interaction of both PNA treatment and age on voltage-dependent activation of total Ca^2+^ conductance (Figure 3A, top, three-way ANOVA, treatment x voltage p=0.0102, age x voltage p=0.0053, Table 4). *Post hoc* analysis revealed differences from -20 to 0mV (3wk VEH vs adult VEH p<0.01 at -20, -10, 0mV; 3wk VEH vs 3wk PNA p<0.05 at -10, 0mV). The potential at which the total Ca^2+^ conductance reaches the half maximum (V_0.5 act_) was depolarized in cells from adult VEH mice compared with those from 3wk VEH mice (Figure 3A, middle, two-way ANOVA age p=0.0304, Fisher’s LSD 3wk VEH vs adult VEH p<0.05, Table 5); this change, however, was not observed in PNA groups. The developmental shift of total Ca^2+^ conductance in VEH mice was associated with a change in medium/slow Ca^2+^ activation (Figure 3B, top, three-way ANOVA age×voltage p<0.0001). *Post hoc* analysis revealed differences from -40 to -20mV (3wk VEH vs adult VEH p<0.01 at -40 and -30mV, p<0.05 at -20mV; 3wk VEH vs 3wk PNA p<0.01 at -40mV, p<0.05 at -30mV). The V_0.5 act_ of slow Ca^2+^ was depolarized in adult VEH group compared with 3wk VEH group (Figure 3B, middle, two-way ANOVA age p=0.0242, Fisher’s LSD 3wk VEH vs adult VEH p<0.05). The V_0.5 act_ of fast Ca^2+^ was unchanged among groups (Figure 3C, middle, two-way ANOVA), although three-way ANOVA of the voltage-dependent fast Ca^2+^ activation curve showed influence of PNA treatment and age (Figure 3C, top, treatment x voltage p=0.0215, age x voltage p=0.0090). The activation slope factor (k) was not influenced by either development or PNA treatment for total Ca^2+^ conductance or the individual components (Figure 3A, B, C, bottom, two-way ANOVA).

**Figure 3.**
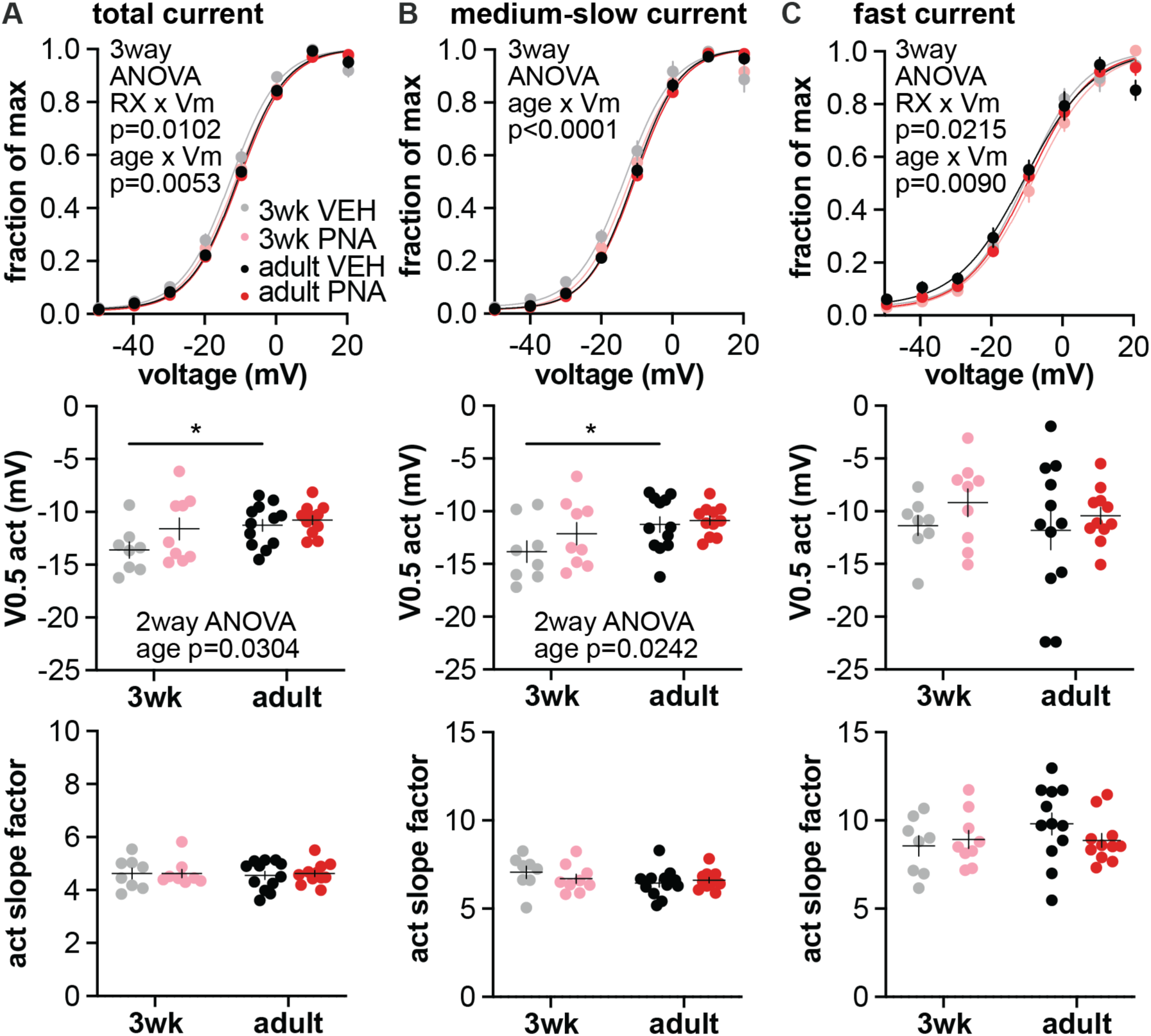
Voltage-dependent activation of Ca^2+^ conductances of GnRH neurons from VEH and PNA mice during development. **A-C** Top: voltage-dependent activation of total (A), medium/slow (B), and fast (C) Ca^2+^ conductance. Dots/error: mean±SEM of Ca^2+^ conductance normalized to the maximum value. Lines: Boltzmann fits of the voltage-dependent activation curves of Ca^2+^ conductance. Middle: half-activation potential (V_0.5 act_) of total (A), medium/slow (B), and fast (C) Ca^2+^ conductance. Bottom: activation slope factor of total (A), medium/slow (B), and fast (C) Ca^2+^ conductance. *p<0.05, Fisher’s LSD. RX, prenatal treatment.

**Table 4.** Three-way ANOVA and Fisher’s LSD test of voltage-dependent activation of calcium conductances (Figure 3). **Bold** indicates p<0.05.

| source of variation | total $I_{Ca}$ | medium/slow $I_{Ca}$ | fast $I_{Ca}$ |
| --- | --- | --- | --- |
| voltage | F (9, 333) = 4282<br>$p < 0.0001$ | F (9, 324) = 2818<br>$p < 0.0001$ | F (9, 324) = 1184<br>$p < 0.0001$ |
| age | F (1, 37) = 3.601<br>$p = 0.0656$ | F (1, 36) = 3.528<br>$p = 0.0685$ | F (1, 36) = 0.2079<br>$p = 0.6512$ |
| treatment | F (1, 37) = 2.547<br>$p = 0.1190$ | F (1, 36) = 2.002<br>$p = 0.1657$ | F (1, 36) = 1.494<br>$p = 0.2295$ |
| voltage x age | F (9, 333) = 2.669<br>$p = 0.0053$ | F (9, 324) = 6.037<br>$p < 0.0001$ | F (9, 324) = 2.205<br>$p = 0.0215$ |
| voltage x treatment | F (9, 333) = 2.454<br>$p = 0.0102$ | F (9, 324) = 1.245<br>$p = 0.2666$ | F (9, 324) = 2.496<br>$p = 0.0090$ |
| age x treatment | F (1, 37) = 0.5269<br>$p = 0.4725$ | F (1, 36) = 1.124<br>$p = 0.2962$ | F (1, 36) = 0.04263<br>$p = 0.8376$ |
| voltage x age x treatment | F (9, 333) = 0.4289<br>$p = 0.9192$ | F (9, 324) = 0.2627<br>$p = 0.9839$ | F (9, 324) = 0.7386<br>$p = 0.6734$ |

| Table 4 continued<br>Post-hoc Fisher's LSD analysis of three-way ANOVA for total calcium conductance activation. |  |  |  |  |
| --- | --- | --- | --- | --- |
| test potential (mV) | 3wk VEH<br>vs.<br>3wk PNA | adult VEH<br>vs.<br>adult PNA | 3wk VEH<br>vs.<br>adult VEH | 3wk PNA<br>vs.<br>adult PNA |
| -50 | $p = 0.6412$ | $p = 0.8314$ | $p = 0.8080$ | $p = 0.9581$ |
| -40 | $p = 0.4119$ | $p = 0.6564$ | $p = 0.6992$ | $p = 0.9452$ |
| -30 | $p = 0.2707$ | $p = 0.5486$ | $p = 0.2970$ | $p = 0.6710$ |
| -20 | $p = 0.1473$ | $p = 0.7388$ | **$p = 0.0036$** | $p = 0.0976$ |
| -10 | **$p = 0.0312$** | $p = 0.4591$ | **$p = 0.0042$** | $p = 0.2123$ |
| 0 | **$p = 0.0223$** | $p = 0.4211$ | **$p = 0.0075$** | $p = 0.3277$ |
| 10 | $p = 0.2275$ | $p = 0.2211$ | $p = 0.7916$ | $p = 0.8968$ |
| 20 | **$p = 0.0146$** | $p = 0.1418$ | $p = 0.1072$ | $p = 0.7062$ |

| Post-hoc Fisher's LSD analysis of three-way ANOVA for medium/slow calcium conductance activation |  |  |  |  |
| --- | --- | --- | --- | --- |
| test potential (mV) | 3wk VEH<br>vs.<br>3wk PNA | adult VEH<br>vs.<br>adult PNA | 3wk VEH<br>vs.<br>adult VEH | 3wk PNA<br>vs.<br>adult PNA |
| -50 | $p = 0.7621$ | $p = 0.8214$ | $p = 0.8073$ | $p = 0.8965$ |
| -40 | $p = 0.3016$ | $p = 0.8934$ | $p = 0.2815$ | $p = 0.9192$ |
| -30 | $p = 0.1341$ | $p = 0.6146$ | $p = 0.0732$ | $p = 0.5038$ |
| -20 | $p = 0.0901$ | $p = 0.9211$ | **$p = 0.0009$** | $p = 0.1414$ |
| -10 | $p = 0.0876$ | **$p = 0.0022$** | $p = 0.1922$ | **$p = 0.0394$** |
| 0 | $p = 0.0976$ | $p = 0.2315$ | $p = 0.0313$ | $p = 0.1314$ |
| 10 | $p = 0.5594$ | $p = 0.6072$ | $p = 0.4311$ | $p = 0.7573$ |
| 20 | $p = 0.2556$ | $p = 0.4045$ | **$p = 0.0012$** | **$p = 0.0044$** |

| Post-hoc Fisher's LSD analysis of three-way ANOVA for fast calcium conductance |  |  |  |  |
| --- | --- | --- | --- | --- |
| test potential (mV) | 3wk VEH<br>vs.<br>3wk PNA | adult VEH<br>vs.<br>adult PNA | 3wk VEH<br>vs.<br>adult VEH | 3wk PNA<br>vs.<br>adult PNA |
| -50 | $p = 0.5930$ | $p = 0.5632$ | $p = 0.8005$ | $p = 0.7660$ |
| -40 | $p = 0.6299$ | $p = 0.2702$ | $p = 0.3478$ | $p = 0.6527$ |
| -30 | $p = 0.6719$ | $p = 0.3713$ | $p = 0.3856$ | $p = 0.6110$ |
| -20 | $p = 0.8388$ | $p = 0.1224$ | $p = 0.2767$ | $p = 0.5815$ |
| -10 | **$p = 0.0322$** | $p = 0.4688$ | $p = 0.9447$ | $p = 0.1150$ |
| 0 | **$p = 0.0172$** | $p = 0.5003$ | $p = 0.4251$ | $p = 0.2512$ |
| 10 | $p = 0.7500$ | $p = 0.4140$ | $p = 0.1633$ | $p = 0.3161$ |
| 20 | $p = 0.1406$ | **$p = 0.0098$** | **$p = 0.0109$** | $p = 0.0751$ |

**Table 5.** Two-way ANOVA of activation properties (Figure 3).

| current type | age | treatment |  | interaction |
| --- | --- | --- | --- | --- |
| total I<sub>Ca</sub> | F (1, 36) = 5.081<br>p=0.0304 | F (1, 36) = 3.155<br>p=0.0842 |  | F (1, 36) = 1.123<br>p=0.2964 |
| medium/slow I<sub>Ca</sub> | F (1, 36) = 5.539<br>p=0.0242 | F (1, 36) = 1.624<br>p=0.2106 |  | F (1, 36) = 0.6619<br>p=0.4212 |
| fast I<sub>Ca</sub> | F (1, 36) = 0.3590<br>p=0.5528 | F (1, 36) = 1.595<br>p=0.2147 |  | F (1, 36) = 0.07312<br>p=0.7884 |
| Fisher's LSD analysis of significant two-way ANOVA |  |  |  |  |
|  | 3wk VEH<br>vs.<br>3wk PNA | adult VEH<br>vs.<br>adult PNA | 3wk VEH<br>vs.<br>adult VEH | 3wk PNA<br>vs.<br>adult PNA |
| total I<sub>Ca</sub> | p=0.0697 | p=0.5858 | p=0.0258 | p=0.4003 |
| medium/slow I<sub>Ca</sub> | p=0.1772 | p=0.7256 | p=0.0326 | p=0.2798 |

**Slope factor**
| current type | age | treatment | interaction |
| --- | --- | --- | --- |
| total I<sub>Ca</sub> | F (1, 36) = 0.05201<br>p=0.8209 | F (1, 36) = 0.04075<br>p=0.8412 | F (1, 36) = 0.04636<br>p=0.8307 |
| medium/slow I<sub>Ca</sub> | F (1, 36) = 1.981<br>p=0.1678 | F (1, 36) = 0.1477<br>p=0.7030 | F (1, 36) = 1.147<br>p=0.2913 |
| fast I<sub>Ca</sub> | F (1, 36) = 1.190<br>p=0.2826 | F (1, 36) = 0.2757<br>p=0.6028 | F (1, 36) = 1.398<br>p=0.2448 |

### Voltage- and time-dependent inactivation, and time-dependent recovery of fast I_Ca_ of GnRH neurons

Representative traces of voltage- and time-dependent inactivation, and time-dependent recovery from inactivation of fast I_Ca_ are in Figures 4A, E, H. PNA treatment shifted the voltage-dependent inactivation curve of GnRH neurons in both 3wk and adult groups to the right (Figure 4B, three-way ANOVA, PNA treatment p=0.0152, PNA×voltage p=0.0106, Table 6). Accordingly, half inactivation potential (V_0.5 inact_) was depolarized in PNA groups, suggesting resistance to voltage-dependent inactivation (Figure 4C, two-way ANOVA, PNA treatment p=0.0169, Table 7). The voltage-dependent inactivation slope factor was not different among groups (Figure 4D, two-way ANOVA). For time-dependent inactivation of the fast I_Ca_, GnRH neurons from 3wk-old mice exhibited a faster inactivation compared to cells from adult groups regardless of PNA treatment (Figure 4F, three-way ANOVA age p=0.0015, age x inactivation time p=0.0040). Accordingly, half inactivation time (T_0.5 inact_) was lower in 3wk-old groups (Figure 4G, two-way ANOVA age p=0.0014, Fisher’s LSD 3wk vs adult p<0.05 in both VEH and PNA groups). Neither age nor PNA treatment altered time-dependent recovery of the fast I_Ca_ (Figure 4I, three-way ANOVA, Figure 4J, two-way ANOVA).

**Figure 4.**
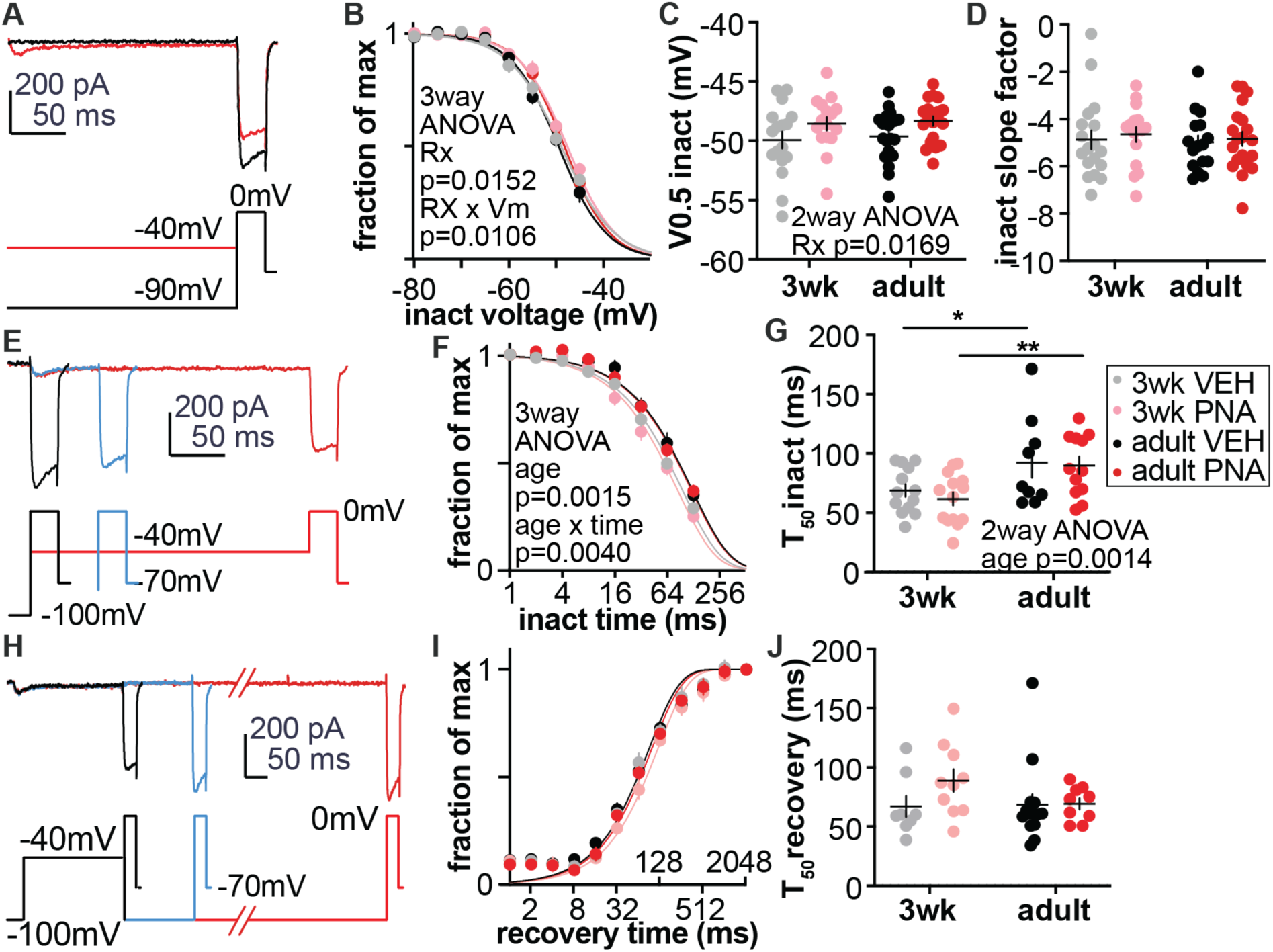
Voltage- and time-dependent inactivation, and time-dependent recovery of GnRH neuron fast I_Ca_ in VEH and PNA mice. **A,** representative current traces (above) and voltage commands (below) for voltage-dependent inactivation of fast I_Ca_. Only two prepulses (-90 and - 40mV) are shown for clarity. **B**, dots/error: mean±SEM of fast I_Ca_ tested at 0mV with different inactivation voltage prepulses. Data were normalized to the maximum value, which is the average of fast I_Ca_ with prepulses of -80, -75 and -70mV. Lines: Boltzmann fits of the voltage-dependent inactivation curves of fast I_Ca_. **C,** half-inactivation potential (V_0.5 inact_) of fast I_Ca_. **D,** inactivation slope factor (k) of fast I_Ca_. **E**, representative current traces (above) and voltage commands (below) for time-dependent inactivation of fast I_Ca_. Only three inactivation times (1, 64, 256ms) are shown for clarity. **F,** dots/error: mean±SEM of fast I_Ca_ tested at 0mV with different inactivation time. Data were normalized to the maximum value obtained with 1ms inactivation time. Lines: monophasic exponential decay fits of time-dependent inactivation curve. **G,** half-inactivation time (T_0.5 inact_) of fast I_Ca_. **H,** representative current traces (above) and voltage commands (below) for time-dependent recovery of fast I_Ca_. Only three recovery times (1, 128, 2048ms) are shown for clarity. **I,** dots/error: mean±SEM of fast I_Ca_ tested at 0mV with different recovery time. Data were normalized to the maximum value obtained after 2048ms recovery. Lines: monophasic exponential association fits of time-dependent recovery curve. **J,** half recovery time (T_0.5 recov_) of fast I_Ca_. *p<0.05, **p<0.01, Fisher’s LSD. RX, prenatal treatment.

**Table 6.** Three-way ANOVA and Fisher’s LSD test of fast component inactivation (Figure 4). **Bold** indicates p<0.05.

| source of variation | voltage-dependent inactivation |  |  |  |
| --- | --- | --- | --- | --- |
| voltage | **F (7, 323) = 540.3, p<0.0001** |  |  |  |
| age | F (1, 69) = 0.1328, p=0.7167 |  |  |  |
| treatment | **F (1, 69) = 6.196, p=0.0152** |  |  |  |
| voltage x age | F (7, 323) = 0.8191, p=0.5719 |  |  |  |
| voltage x treatment | **F (7, 323) = 2.673, p=0.0106** |  |  |  |
| age x treatment | F (1, 69) = 0.1502, p=0.6996 |  |  |  |
| voltage x age x treatment | F (7, 323) = 0.2909, p=0.9573 |  |  |  |
| Fisher's LSD analysis of three-way ANOVA for voltage-dependent inactivation of the fast component |  |  |  |  |
| **inactivation prepulse (mV)** | 3wk VEH<br>vs.<br>3wk PNA | adult VEH<br>vs.<br>adult PNA | 3wk VEH<br>vs.<br>adult VEH | 3wk PNA<br>vs.<br>adult PNA |
| -80 | p=0.7487 | p=0.6050 | p=0.9253 | p=0.9507 |
| -75 | p=0.8485 | p=0.7855 | p=0.5042 | p=0.5696 |
| -70 | p=0.7752 | p=0.9512 | p=0.7249 | p=0.9210 |
| -65 | p=0.2767 | p=0.8134 | p=0.5259 | p=0.8301 |
| -60 | **p=0.0432** | p=0.6674 | p=0.1975 | p=0.6530 |
| -55 | **p=0.0131** | **p=0.0118** | p=0.5156 | p=0.5733 |
| -50 | **p=0.0409** | **p=0.0285** | p=0.8807 | p=0.8365 |
| -45 | p=0.2268 | p=0.3283 | p=0.2852 | p=0.2088 |

**time-dependent inactivation and recovery**
| source of variation | time-dependent inactivation | time-dependent recovery |  |  |
| --- | --- | --- | --- | --- |
| time | **F (7, 301) = 572.2**<br>**p<0.0001** | **F (11, 407) = 1125**<br>**p<0.0001** |  |  |
| age | **F (1, 43) = 11.52**<br>**p=0.0015** | F (1, 37) = 0.4679<br>p=0.4982 |  |  |
| treatment | F (1, 43) = 0.3162<br>p=0.5768 | F (1, 37) = 2.367<br>p=0.1324 |  |  |
| time x age | **F (7, 301) = 3.061**<br>**p=0.0040** | F (11, 407) = 0.4952<br>p=0.9062 |  |  |
| time x treatment | F (7, 301) = 1.481<br>p=0.1734 | F (11, 407) = 0.8101<br>p=0.6300 |  |  |
| age x treatment | F (1, 43) = 0.09984<br>p=0.7536 | F (1, 37) = 0.3475<br>p=0.5591 |  |  |
| time x age x treatment | F (7, 301) = 0.5131<br>p=0.8245 | F (11, 407) = 1.191<br>p=0.2909 |  |  |
| Fisher's LSD analysis of three-way ANOVA time dependent inactivation of the fast component |  |  |  |  |
| **duration of inactivation (s)** | 3wk VEH<br>vs.<br>3wk PNA | adult VEH<br>vs.<br>adult PNA | 3wk VEH<br>vs.<br>adult VEH | 3wk PNA<br>vs.<br>adult PNA |
| 1 | p>0.9999 | p>0.9999 | p>0.9999 | p>0.9999 |
| 2 | p=0.4712 | p=0.7016 | p=0.7076 | p=0.8869 |
| 4 | p=0.4776 | p=0.9301 | p=0.2109 | p=0.4434 |
| 8 | p=0.5375 | p=0.8251 | p=0.2042 | p=0.3057 |
| 16 | p=0.0552 | p=0.2665 | **p=0.0469** | **p=0.0064** |
| 32 | p=0.0871 | p=0.9330 | p=0.1045 | **p=0.0011** |
| 64 | p=0.5163 | p=0.4257 | **p=0.0150** | **p=0.0187** |
| 128 | p=0.2507 | p=0.6265 | p=0.1135 | **p=0.0009** |

**Table 7.** Two-way ANOVA of inactivation properties (Figure 4).

| property | age |  | treatment |  | interaction |
| --- | --- | --- | --- | --- | --- |
| V<sub>50 inact</sub> | F (1, 68) = 0.2172<br>p=0.6426 |  | F (1, 68) = 5.993<br>p=0.0169 |  | F (1, 68) = 0.005608<br>p=0.9405 |
| voltage-dependent inactivation slope factor | F (1, 66) = 0.1908<br>p=0.6637 |  | F (1, 66) = 0.3048<br>p=0.5827 |  | F (1, 66) = 0.01175<br>p=0.9140 |
| T<sub>50 inact</sub> | F (1, 44) = 11.63<br>p=0.0014 |  | F (1, 44) = 0.3672<br>p=0.5476 |  | F (1, 44) = 0.1003<br>p=0.7529 |
| T<sub>50 recov</sub> | F (1, 37) = 0.9851<br>p=0.3274 |  | F (1, 37) = 1.565<br>p=0.2188 |  | F (1, 37) = 1.337<br>p=0.2551 |
| Fisher's LSD analysis of significant two-way ANOVA |  |  |  |  |  |
|  | 3wk VEH<br>vs.<br>3wk PNA | adult VEH<br>vs.<br>adult PNA | 3wk VEH<br>vs.<br>adult VEH | 3wk PNA<br>vs.<br>adult PNA |  |
| V<sub>50 inact</sub> | p=0.0911 | p=0.0841 | p=0.7026 | p=0.7834 |  |
| T<sub>50 inact</sub> | p=0.4868 | p=0.8486 | p=0.0426 | p=0.0081 |  |

### Excitability of GnRH neurons is affected by development in VEH mice

The excitability of GnRH neurons was quantified as the number of action potentials generated in response to 500ms current injections during whole-cell current-clamp recordings. In pilot experiments, we used EGTA-containing vs. EGTA-free pipette solution to assess the effects of exogenous Ca^2+^ buffering on GnRH neuron action potential firing and membrane potential trajectory after current injection. Fewer spikes were generated in response to current injections when GnRH neurons were recorded with EGTA-free compared to those recorded using EGTA-containing pipette solution (two-way ANOVA current F(6, 42)=38.82, p<0.0001; EGTA F(1, 7)=6.362, p=0.0397; current x EGTA F(6, 42) p<0.0001). A post-spike-train membrane hyperpolarization was observed when EGTA-free solution was used (Figure 5A bottom, arrow), which was not observed in recordings using EGTA-containing solution (Figure 5A top). We thus used EGTA-free pipette solution in recordings to minimize effects of buffering on neuron excitability. GnRH neuron capacitance was higher in adult compared to 3wk-old groups (p=0.0317), while PNA treatment had no effect; R_in_, R_s_, and I_hold_ were not different among groups (Table 8, two-way ANOVA). Three-way ANOVA of firing rate in response to current injection steps revealed an effect of age on GnRH neurons excitability (figure 5B, Table 9 age x current p=0.0034). The difference was driven by higher excitability in cells from 3wk-old VEH compared to adult VEH mice (Fisher’s LSD 3wk VEH vs adult VEH p<0.05 at 50pA, p<0.01 at 60 to 90pA, p<0.001 at 100pA). In contrast, excitability of GnRH neurons from PNA mice showed no difference with development. Two-way ANOVA of firing rate in response to 100pA injection confirmed increased firing rate in cells from VEH but not PNA mice at 3wks-of-age, with no difference between VEH and PNA groups at either age (Figure 5C, age F(1, 58)=4.418, p=0.0399; PNA treatment F(1, 58)=0.1243, p=0.7257; interaction F(1, 58)=1.945, p=0.1685; Fisher’s LSD 3wk VEH vs adult VEH p<0.05).

**Figure 5.**
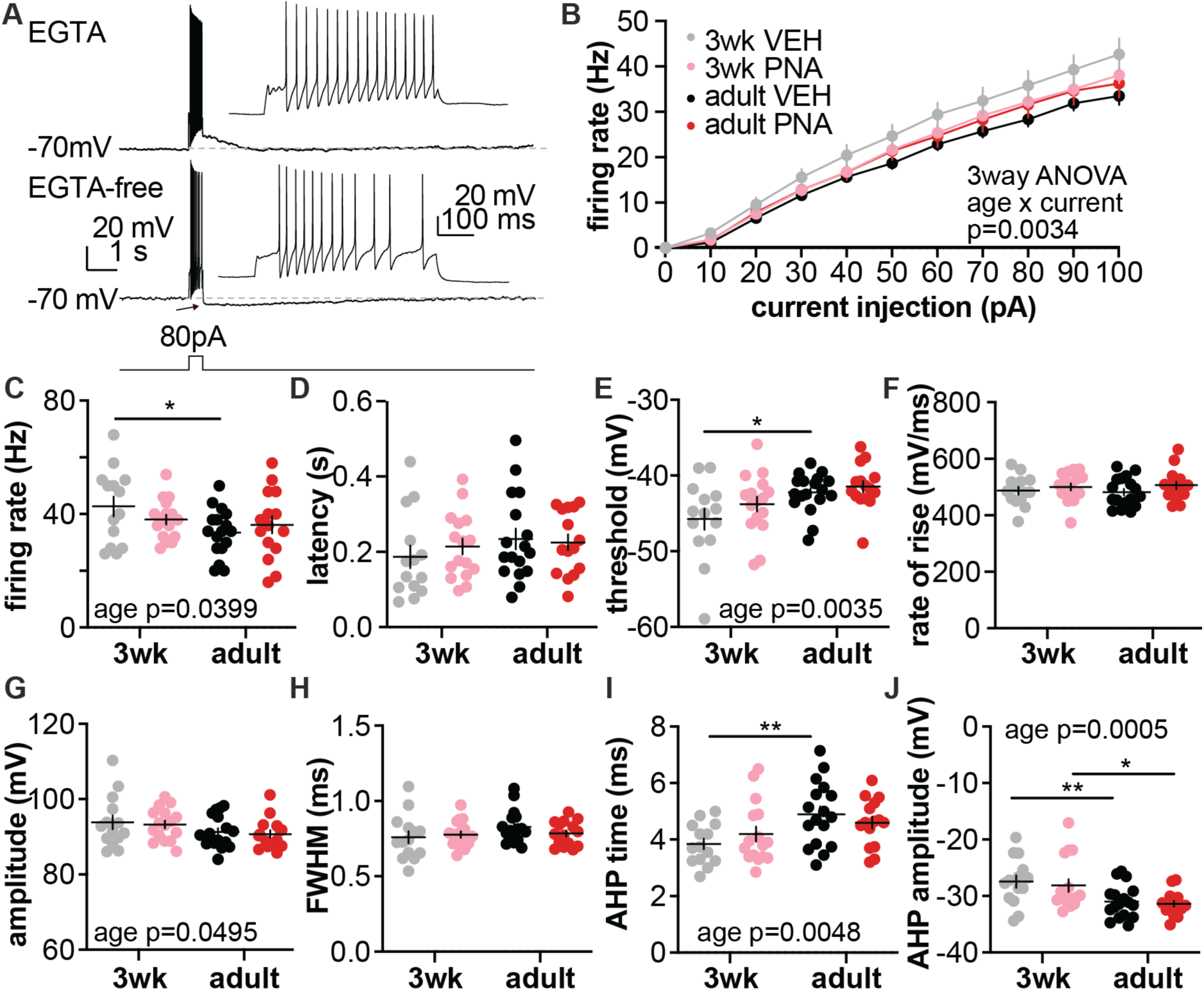
Intrinsic excitability of GnRH neurons in VEH and PNA mice during development. **A,** representative recordings of a GnRH neuron membrane potential in response to 80pA current injection using EGTA-containing (top) vs EGTA-free pipette solution (bottom, used for the rest of the recordings, arrow shows post-spike-train hyperpolarization). Insets show individual action potentials. **B,** mean±SEM firing rate during current injections from 10 to 100pA with EGTA-free solutions. **C,** Firing rate during 100pA current injection. **D-J,** the first action potential latency (D), threshold (E), rise rate (F), amplitude relative to threshold (G), full-width at half maximum (H), AHP time (I), AHP amplitude relative to threshold. *p<0.05, **p<0.01, Fisher’s LSD.

**Table 8.** descriptive statistics and two-way ANOVA parameters for passive properties and recording quality for GnRH neuron excitability recording (Figure 5). **Bold** indicates p<0.05.

| Descriptive statistics |  |  |  |  |
| --- | --- | --- | --- | --- |
| property | 3wk VEH | 3wk PNA | adult VEH | adult PNA |
| series resistance (MΩ) | 16.594 ± 0.770 | 16.427 ± 0.717 | 15.511 ± 1.062 | 14.833 ± 0.554 |
| input resistance (MΩ) | 1280.424 ± 100.633 | 1098.038 ± 75.181 | 1192.588 ± 62.423 | 1222.051 ± 79.011 |
| capacitance (pF) | 12.739 ± 0.689 | 13.202 ± 0.623 | 15.045 ± 0.568 | 14.371 ± 0.899 |
| holding current (pA) | -6.327 ± 2.300 | -7.228 ± 2.179 | -5.550 ± 1.295 | -5.752 ± 1.956 |
| Two-way ANOVA |  |  |  |  |
|  | age | treatment | interaction |  |
| series resistance (MΩ) | F(1,58)= 2.660<br>p=0.1083 | F(1,58)= 0.2646<br>p=0.6090 | F(1, 58)= 0.09678<br>p=0.7568 |  |
| input resistance (MΩ) | F(1,58)= 0.05253<br>p=0.8195 | F(1,58)= 0.9386<br>p=0.3367 | F(1,58)= 1.801<br>p=0.1848 |  |
| capacitance (pF) | **F(1,58)= 6.175**<br>**p=0.0159** | F(1,58)= 0.02271<br>p=0.8807 | F(1,58)= 0.4195<br>p=0.2669 |  |
| holding current (pA) | F(1,58)= 0.5583<br>p=0.4579 | F(1,58)= 0.2039<br>p=0.6533 | F(1,58)= 0.0001689<br>p=0.9897 |  |
| Fisher's LSD analysis of significant two-way ANOVA |  |  |  |  |
|  | 3wk VEH<br>vs.<br>3wk PNA | adult VEH<br>vs.<br>adult PNA | 3wk VEH<br>vs.<br>adult VEH | 3wk PNA<br>vs.<br>adult PNA |
| capacitance (pF) | p=0.6465 | p=0.4911 | **p=0.0235** | p=0.0589 |

**Table 9.** Three-way ANOVA and Fisher’s LSD test of GnRH neuron excitability (Figure 5). **Bold** indicates p<0.05.

| source of variation |  |  |  |  |
| --- | --- | --- | --- | --- |
| voltage | **F (10, 580) = 697.7, p<0.0001** |  |  |  |
| age | F (1, 58) = 3.424, p=0.0693 |  |  |  |
| treatment | F (1, 58) = 0.1387, p=0.7110 |  |  |  |
| voltage x age | **F (10, 580) = 2.667, p=0.0034** |  |  |  |
| voltage x treatment | F (10, 580) = 0.1988, p=0.9964 |  |  |  |
| age x treatment | F (1, 58) = 2.389, p=0.1276 |  |  |  |
| voltage x age x treatment | F (10, 580) = 1.307, p=0.2228 |  |  |  |
| Fisher's LSD analysis of three-way ANOVA for firing rate at different current injections |  |  |  |  |
| current injection (pA) | 3wk VEH vs. 3wk PNA | adult VEH vs. adult PNA | 3wk VEH vs. adult VEH | 3wk PNA vs. adult PNA |
| 0 | p>0.9999 | p>0.9999 | p>0.9999 | p>0.9999 |
| 10 | p=0.5858 | p=0.8241 | p=0.4087 | p=0.9556 |
| 20 | p=0.4236 | p=0.6099 | p=0.2428 | p=0.8853 |
| 30 | p=0.2759 | p=0.6454 | p=0.1245 | p=0.9843 |
| 40 | p=0.1556 | p=0.6841 | p=0.0614 | p=0.9738 |
| 50 | p=0.2329 | p=0.2945 | **p=0.0188** | p=0.9086 |
| 60 | p=0.1177 | p=0.4620 | **p=0.0099** | p=0.7805 |
| 70 | p=0.3009 | p=0.3182 | **p=0.0092** | p=0.5596 |
| 80 | p=0.1637 | p=0.1953 | **p=0.0034** | p=0.7982 |
| 90 | p=0.0981 | p=0.2667 | **p=0.0038** | p=0.8957 |
| 100 | p=0.0766 | p=0.2748 | **p=0.0003** | p=0.4648 |

The properties of the first action potential generated in EGTA-free recordings were analyzed by two-way ANOVA (Figure 5D-J, Table 10). Action potential threshold was hyperpolarized in GnRH neurons from 3wk vs adult VEH mice (Figure 5E, age p=0.0035, Fisher’s LSD 3wk VEH vs adult VEH p<0.05), and the peak of the afterhyperpolarization (AHP) was earlier (Figure 5I, age p=0.0048, Fisher’s LSD 3wk VEH vs adult VEH p<0.01). The amplitude of AHP relative to threshold was lower in cells from 3wk than those from adults in both VEH and PNA groups (Figure 5J, age p=0.0005, Fisher’s LSD 3wk VEH vs adult VEH p<0.01, 3wk PNA vs adult PNA p<0.05). Action potential amplitude relative to threshold was higher in GnRH neurons from 3wk-old mice compared with those from adults (Figure 5G, age p=0.0495). The changes in AHP and action potential amplitude in 3wk-old mice were likely attributed to the hyperpolarized threshold (Figure 5E), as the absolute peak values of AHP and action potential were not different among groups (absolute AHP peak: 3wk VEH -73.2±0.7mV, 3wk PNA -71.9±0.7mV, adult VEH - 73.3±0.7mV, adult PNA -72.8±0.5mV; action potential peak: 3wk VEH 48.1±0.9mV, 3wk PNA 49.5±0.8mV, adult VEH 49.0±0.8mV, adult PNA 49.2±0.7mV, two-way ANOVA). There was no difference in latency to fire in response to current injection, action potential rate of rise, and full width at half maximum (FWHM) (Figure 5D, F, H).

**Table 10.** Two-way ANOVA of first action potential properties (Figure 5).

| property | age | treatment |  | interaction |
| --- | --- | --- | --- | --- |
| latency | F (1, 58) = 1.287<br>p=0.2613 | F (1, 58) = 0.1300<br>p=0.7197 |  | F (1, 58) = 0.5075<br>p=0.4791 |
| threshold | F (1, 59) = 9.253<br>p=0.0035 | F (1, 59) = 2.082<br>p=0.1543 |  | F (1, 59) = 0.4062<br>p=0.5263 |
| rate of rise | F (1, 58) = 0.0003850<br>p=0.9844 | F (1, 58) = 2.020<br>p=0.1606 |  | F (1, 58) = 0.2035<br>p=0.6536 |
| AP amplitude | F (1, 58) = 4.026<br>p=0.0495 | F (1, 58) = 0.2041<br>p=0.6531 |  | F (1, 58) = 1.866e-006<br>p=0.9989 |
| FWHM | F (1, 58) = 1.864<br>p=0.1774 | F (1, 58) = 0.2212<br>p=0.6399 |  | F (1, 58) = 1.195<br>p=0.2789 |
| AHP time | F (1, 58) = 8.584<br>p=0.0048 | F (1, 58) = 0.01039<br>p=0.9192 |  | F (1, 58) = 1.747<br>p=0.1914 |
| AHP amplitude | F (1, 58) = 13.78<br>p=0.0005 | F (1, 58) = 0.3468<br>p=0.5582 |  | F (1, 58) = 0.02400<br>p=0.8774 |
| Fisher's LSD analysis of significant two-way ANOVA |  |  |  |  |
|  | 3wk VEH<br>vs.<br>3wk PNA | adult VEH<br>vs.<br>adult PNA | 3wk VEH<br>vs.<br>adult VEH | 3wk PNA<br>vs.<br>adult PNA |
| threshold | p=0.1556 | p=0.5620 | **p=0.0114** | p=0.0957 |
| amplitude | p=0.7536 | p=0.7472 | p=0.1619 | p=0.1607 |
| AHP time | p=0.3259 | p=0.3841 | **p=0.0040** | p=0.2592 |
| AHP amplitude | p=0.6067 | p=0.7561 | **p=0.0084** | **p=0.0145** |

### Blocking small-conductance Ca²⁺-activated K⁺ (SK) current increases GnRH neuron firing except in adult PNA mice

Calcium influx and [Ca^2+^]_in_ could activate SK channels, leading to a hyperpolarizing SK current. We tested the role of the SK current on GnRH neuron intrinsic excitability by blocking it with apamin (Figure 6A, B). As in the other experiments, GnRH neuron capacitance was higher in adult compared to 3wk-old groups, while PNA treatment had no effect; R_in_, R_s_, and I_hold_ were not different among groups (Table 11, two-way ANOVA).GnRH neuron firing response to a train of current injections (20, 40, 80pA) was recorded under basal conditions (0min) and 2, 4, and 6min after the onset of apamin or control perfusion (Figure 6C-N). During 6-min recording, ACSF control cells from 3wk-old VEH and PNA mice maintained a relative consistent firing in response to the subsequent same current injections (Figure 6C, D, G, H, K, L, black trace). ACSF control cells from adult VEH and PNA mice exhibited a progressive decrease in firing frequency (Figure 6E, F, I, J, M, N, black trace). Blocking the SK current with apamin progressively increased the 80pA-triggered GnRH neuron firing rate in cells from both 3wk-old and adult VEH mice (Figure 6C, E, two-way ANOVA time p<0.05, apamin p<0.05, time x apamin p<0.0001 for both, Table 12), with higher firing rates observed 4 and 6min after onset of apamin treatment compared with 0min basal state (Fisher’s LSD p<0.001 for both) and the ACSF controls (Fisher’s LSD p<0.01 for both). In GnRH neurons from 3wk-old PNA mice, apamin treatment led to an increase of 80pA-triggered GnRH neuron firing rate compared with the 0min basal state (Figure 6D, two-way ANOVA, time x apamin p<0.0001, Fisher’s LSD 6min vs 0min p<0.0001) and the ACSF controls (Fisher’s LSD p<0.05 at 4 and 6min). In GnRH neurons from PNA adults, apamin perfusion mitigated the time-dependent loss of GnRH neuron firing, evident as higher firing rate at 6min after the onset of apamin treatment compared with the ACSF controls (Figure 6F, two-way ANOVA time x apamin p=0.0221, Fisher’s LSD p<0.05). The firing rate, however, did not increase further as observed in other groups and remained comparable to the 0min basal level (Fisher’s LSD), indicating a blunted response to apamin in PNA adults. Response to 40pA current injections was less robust in terms of firing rate, as expected, but all groups showed a similar response to apamin as observed during 80pA current injection (Figure 6G-J). GnRH neurons had minimal firing in response to 20pA current injections and the effect apamin was less pronounced (Figure 6K-N). We thus focused on the data obtained at 80pA current injection in the following analysis.

**Figure 6.**
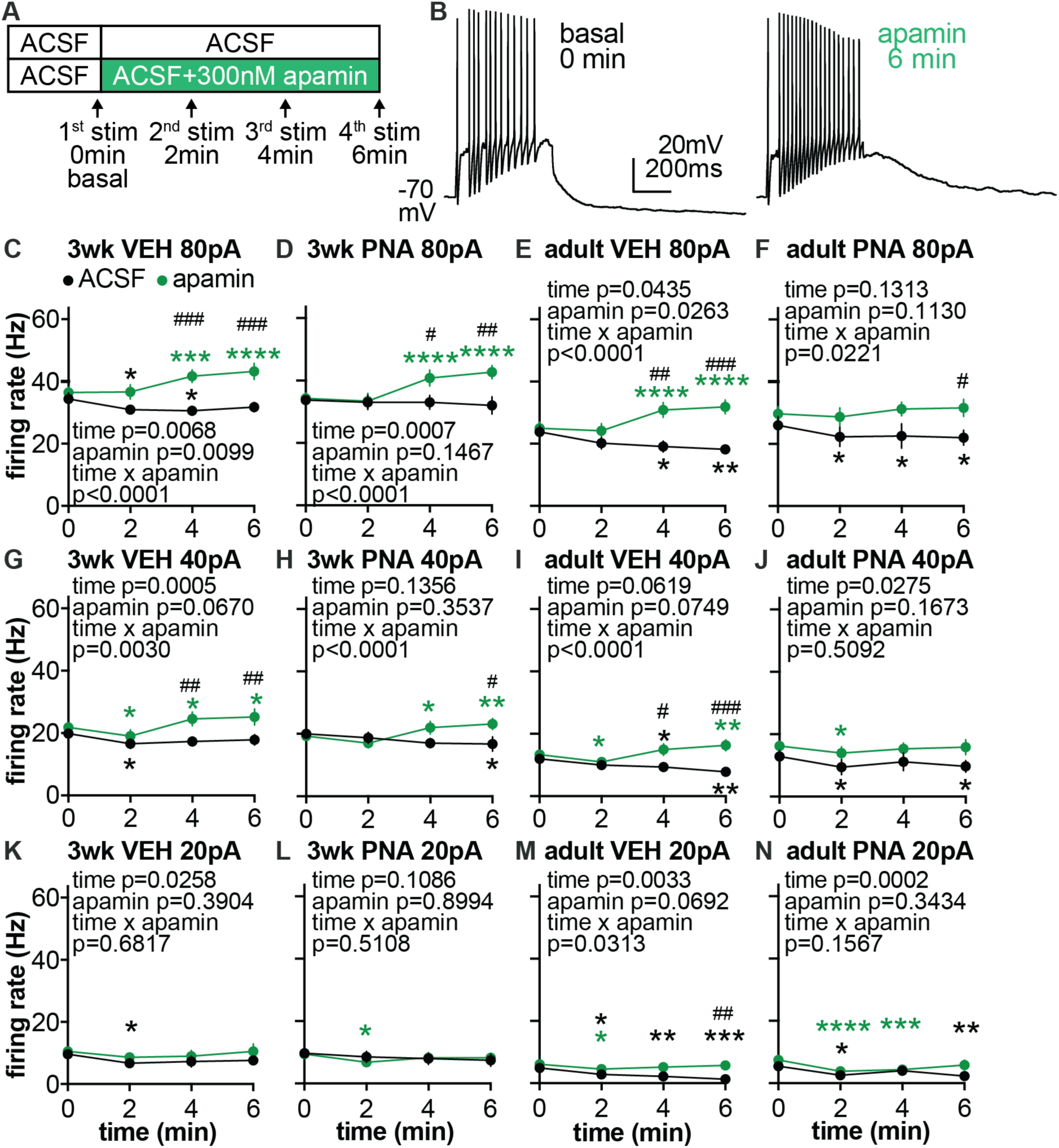
Effect of apamin perfusion on the firing rate of GnRH neurons from 3wk-old and adult VEH and PNA mice. **A,** timeline of current injection trains (20, 40, 80pA). **B,** representative action potential firing of a GnRH neuron from a 3wk-VEH mouse in response to 80pA current injection at 0min basal state and at 6min of apamin perfusion. **C-N,** Firing rate during current injection under 0min basal state and at 2, 4, and 6min of apamin (green) or ACSF (black) perfusion. *p<0.05, **p<0.01, ***p<0.001 compared with 0min basal state within the group indicated by the corresponding color code; ^#^p<0.05, ^##^p<0.01, ^###^p<0.001 apamin compared with ACSF control, Fisher’s LSD.

**Table 11.** descriptive statistics and two-way ANOVA parameters for passive properties and recording quality for GnRH neuron excitability recording with vehicle or apamin perfusion (Figure 6, 7, 8). **Bold** indicates p<0.05.

| Descriptive statistics |  |  |  |  |
| --- | --- | --- | --- | --- |
| property | 3wk VEH | 3wk PNA | adult VEH | adult PNA |
| series resistance (MΩ) | 17.115 ± 0.545 | 16.168 ± 0.757 | 16.534 ± 0.647 | 16.895 ± 0.708 |
| input resistance (MΩ) | 1527.087±<br>107.921 | 1507.790 ± 94.474 | 1417.696 ± 96.553 | 1364.739 ± 84.815 |
| capacitance (pF) | 12.498 ± 0.402 | 13.246 ± 0.366 | 16.003 ± 0.426 | 15.091± 0.617 |
| holding current (pA) | -6.465 ± 1.470 | -2.064 ± 1.040 | -2.048 ± 1.334 | -2.734 ± 1.785 |
| Two-way ANOVA |  |  |  |  |
|  | age | treatment | interaction |  |
| series resistance (MΩ) | F(1,86)= 0.1165<br>p=0.9143 | F(1,86)= 0.1885<br>p=0.6652 | F(1, 86)= 0.9390<br>p=0.3353 |  |
| input resistance (MΩ) | F(1,86)= 1.673<br>p=0.1993 | F(1,86)= 0.1371<br>p=0.7121 | F(1,86)= 0.02975<br>p=0.8635 |  |
| capacitance (pF) | **F(1,86)= 33.26**<br>**P<0.0001** | F(1,86)= 0.03091<br>p=0.8609 | F(1,86)= 3.206<br>p=0.0769 |  |
| holding current (pA) | F(1,86)= 1.700<br>p=0.1958 | F(1,86)= 1.670<br>p=0.1998 | F(1,86)= 3.131<br>p=0.0804 |  |
| Fisher's LSD analysis of significant two-way ANOVA |  |  |  |  |
|  | 3wk VEH<br>vs.<br>3wk PNA | adult VEH<br>vs.<br>adult PNA | 3wk VEH<br>vs.<br>adult VEH | 3wk PNA<br>vs.<br>adult PNA |
| capacitance (pF) | p=0.6847 | p=0.4824 | **p<0.0001** | **p=0.0375** |

**Table 12.** Two-way ANOVA of the effects of apamin (Figure 6). **Blue:** decrease compared with 0min baseline; red: increased compared with 0min baseline.

| group | time |  | apamin |  | interaction |  |  |
| --- | --- | --- | --- | --- | --- | --- | --- |
| 80pA 3wk VEH | F (3, 63) = 4.435<br>p=0.0068 |  | F (1, 21) = 8.038<br>p=0.0099 |  | F (3, 63) = 9.540<br>p<0.0001 |  |  |
| 80pA 3wk PNA | F (3, 60) = 6.444<br>p=0.0007 |  | F (1, 20) = 2.280<br>p=0.1467 |  | F (3, 60) = 10.23<br>p<0.0001 |  |  |
| 80pA adult VEH | F (3, 75) = 2.842<br>p=0.0435 |  | F (1, 25) = 5.580<br>p=0.0263 |  | F (3, 75) = 14.42<br>p<0.0001 |  |  |
| 80pA adult PNA | F (3, 60) = 1.949<br>p=0.1313 |  | F (1, 20) = 2.748<br>p=0.1130 |  | F (3, 60) = 3.446<br>p=0.0221 |  |  |
| Fisher's LSD analysis of significant two-way ANOVA |  |  |  |  |  |  |  |
|  | ACSF |  |  |  |  |  |  |
| comparison | 3wk VEH |  | 3wk PNA |  | adult VEH |  | adult PNA |
| 0min vs 2min | p=0.0329 |  | p=0.6850 |  | p=0.0549 |  | p=0.0273 |
| 0min vs 4min | p=0.0176 |  | p=0.6850 |  | p=0.0125 |  | p=0.0410 |
| 0min vs 6min | p=0.1002 |  | p=0.3453 |  | p=0.0032 |  | p=0.0178 |
|  | apamin |  |  |  |  |  |  |
|  | 3wk VEH |  | 3wk PNA |  | adult VEH |  | adult PNA |
| 0min vs 2min | p=0.9107 |  | p=0.5400 |  | p=0.5482 |  | p=0.3454 |
| 0min vs 4min | p=0.0003 |  | p<0.0001 |  | p<0.0001 |  | p=0.1960 |
| 0min vs 6min | p<0.0001 |  | p<0.0001 |  | p<0.0001 |  | p=0.1013 |
|  | ACSF vs apamin |  |  |  |  |  |  |
|  | 3wk VEH |  | 3wk PNA |  | adult VEH |  | adult PNA |
| 0min | p=0.4866 |  | p=0.8797 |  | p=0.7293 |  | p=0.4034 |
| 2min | p=0.0604 |  | p=0.9036 |  | p=0.2588 |  | p=0.1550 |
| 4min | p=0.0003 |  | p=0.0288 |  | p=0.0012 |  | p=0.0557 |
| 6min | p=0.0002 |  | p=0.0032 |  | p=0.0002 |  | p=0.0338 |

| group | time | apamin | interaction |  |
| --- | --- | --- | --- | --- |
| 40pA 3wk VEH | F (3, 63) = 6.784<br>p=0.0005 | F (1, 21) = 3.732<br>p=0.0670 | F (3, 63) = 5.166<br>p=0.0030 |  |
| 40pA 3wk PNA | F (3, 60) = 1.922<br>p=0.1356 | F (1, 20) = 0.9017<br>p=0.3537 | F (3, 60) = 9.179<br>p<0.0001 |  |
| 40pA adult VEH | F (3, 75) = 2.552<br>p=0.0619 | F (1, 25) = 3.455<br>p=0.0749 | F (3, 75) = 9.590<br>p<0.0001 |  |
| 40pA adult PNA | F (3, 60) = 3.261<br>p=0.0275 | F (1, 20) = 2.053<br>p=0.1673 | F (3, 60) = 0.7809<br>p=0.5092 |  |
| Fisher's LSD analysis of significant two-way ANOVA |  |  |  |  |
|  | ACSF |  |  |  |
| comparison | 3wk VEH | 3wk PNA | adult VEH | adult PNA |
| 0min vs 2min | p=0.0154 | p=0.4119 | p=0.1439 | p=0.0282 |
| 0min vs 4min | p=0.0572 | p=0.0519 | p=0.0526 | p=0.2654 |
| 0min vs 6min | p=0.1329 | p=0.0357 | p=0.0026 | p=0.0436 |
|  | apamin |  |  |  |
|  | 3wk VEH | 3wk PNA | adult VEH | adult PNA |
| 0min vs 2min | p=0.0278 | p=0.0502 | p=0.0172 | p=0.0335 |
| 0min vs 4min | p=0.0380 | p=0.0208 | p=0.0858 | p=0.3738 |
| 0min vs 6min | p=0.0102 | p=0.0013 | p=0.0025 | p=0.7023 |
|  | ACSF vs apamin |  |  |  |
|  | 3wk VEH | 3wk PNA | adult VEH | adult PNA |
| 0min | p=0.4583 | p=0.7860 | p=0.5871 | p=0.3177 |
| 2min | p=0.3667 | p=0.5064 | p=0.6837 | p=0.1814 |
| 4min | p=0.0091 | p=0.0601 | p=0.0226 | p=0.2202 |
| 6min | p=0.0080 | p=0.0170 | p=0.0007 | p=0.0734 |
| 20pA 3wk VEH | F (3, 63) = 3.305<br>p=0.0258 | F (1, 21) = 0.7693<br>p=0.3904 | F (3, 63) = 0.5029<br>p=0.6817 |  |
| 20pA 3wk PNA | F (3, 60) = 2.108<br>p=0.1086 | F (1, 20) = 0.01639<br>p=0.8994 | F (3, 60) = 0.7781<br>p=0.5108 |  |
| 20pA adult VEH | F (3, 75) = 4.991<br>p=0.0033 | F (1, 25) = 3.606<br>p=0.0692 | F (3, 75) = 3.110<br>p=0.0313 |  |
| 20pA adult PNA | F (3, 60) = 7.550<br>p=0.0002 | F (1, 20) = 0.9420<br>p=0.3434 | F (3, 60) = 1.801<br>p=0.1567 |  |
| Fisher's LSD analysis of significant two-way ANOVA |  |  |  |  |
|  | ACSF |  |  |  |
| comparison | 3wk VEH | 3wk PNA | adult VEH | adult PNA |
| 0min vs 2min | 0.0167 |  | 0.0388 | 0.0177 |
| 0min vs 4min | 0.0500 |  | 0.0064 | 0.2276 |
| 0min vs 6min | 0.0958 |  | 0.0004 | 0.0094 |
|  | apamin |  |  |  |
|  | 3wk VEH | 3wk PNA | adult VEH | adult PNA |
| 0min vs 2min | 0.1105 |  | 0.0235 | <0.0001 |
| 0min vs 4min | 0.1902 |  | 0.1903 | 0.0003 |
| 0min vs 6min | >0.9999 |  | 0.6216 | 0.0502 |
|  | ACSF vs apamin |  |  |  |
|  | 3wk VEH | 3wk PNA | adult VEH | adult PNA |
| 0min | 0.7123 |  | 0.4281 | 0.3289 |
| 2min | 0.4084 |  | 0.2805 | 0.5342 |
| 4min | 0.4613 |  | 0.0536 | 0.8476 |
| 6min | 0.2233 |  | 0.0047 | 0.0882 |

### Components of post-spike-train membrane potential changes are altered in GnRH neurons from PNA mice

Under basal conditions (before apamin), at the end of the current injection the trajectory of the membrane potential change exhibited two phases. Immediately after termination of the current injection, most cells rapidly repolarized, with 46 of 61 cells across the four groups showing fast membrane hyperpolarization (fast post-spike-train ΔVm relative to baseline potential, magenta Figure 7A). All cells then exhibited a more prolonged, slow hyperpolarization, which gradually returned to baseline potential over several seconds (slow post-spike-train ΔVm relative to baseline potential, cyan Figure 7A). The area under the curve (AUC) of ΔV_m_ relative to baseline potential was not different among groups during the 12.5s post-stimulus period (Figure 7B, two-way ANOVA, Table 13). To quantify the fast vs slow post-spike-train ΔV_m_, the inflection point when the repolarization rate after current injection termination fell below 50mV/s was used, with the period before being designated fast and that after slow (Figure 7A, arrow). The fast ΔV_m_ relative to baseline potential was shifted towards zero or positive values in cells from PNA mice (Figure 7C, two-way ANOVA, PNA treatment p=0.0390). The latency to reach the inflection point was longer in GnRH neurons from 3wk PNA than adult PNA mice (two-way ANOVA, age p=0.0112, Fisher’s LSD 3wk PNA vs adult PNA p<0.01), suggesting a prolonged repolarization in younger PNA animals (Figure 7E). The peak value and latency time of post-spike-train slow ΔV_m_ were not different among groups (Figure 7D, F).

**Figure 7.**
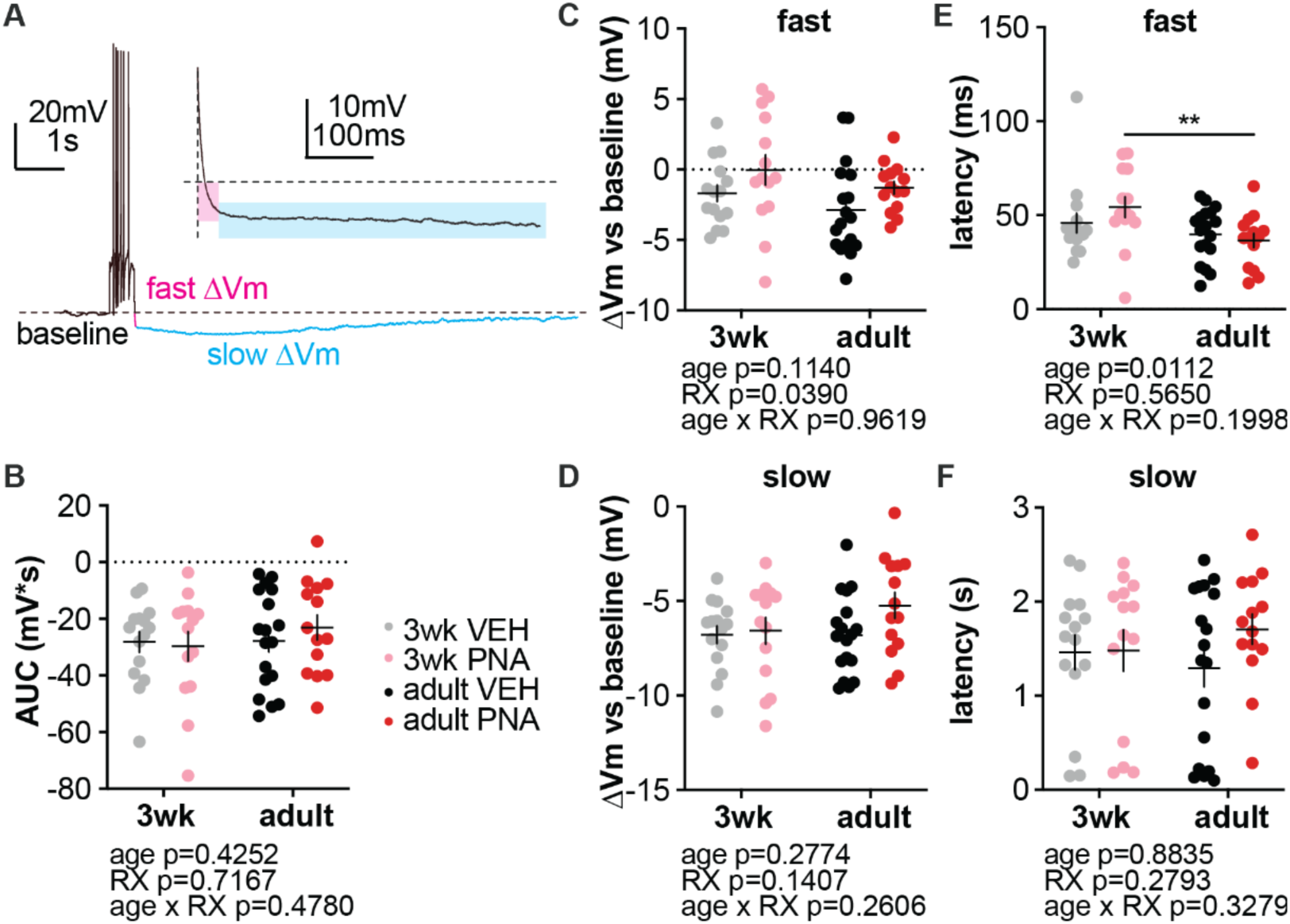
Change in post-spike-train membrane potential (ΔV_m_) relative to baseline potential under basal conditions. **A,** representative cell from an adult VEH mouse in response to 80pA current injection. Dashed line: pre-stimulus baseline; magenta to cyan color change marks repolarization rate falling below 50mV/s. Insert: expanded trace showing fast ΔV_m_ (magenta box) and slow ΔVm (cyan box). **B,** post-spike-train ΔV_m_ area under the curve (AUC) for the 12.5s analysis period**. C, D,** post-spike-train fast ΔV_m_ peak value (C) and latency (D). **E, F,** post-spike-train slow ΔV_m_ peak value (E) and latency (F). RX, prenatal treatment. **p<0.01, Fisher’s LSD.

**Table 13.** Two-way ANOVA of post-spike train change in membrane potential under basal conditions (Figure 7).

| proerty | age | treatment |  | interaction |
| --- | --- | --- | --- | --- |
| $\Delta V_m$ AUC | F (1, 57) = 0.6452<br>p=0.4252 | F (1, 57) = 0.1330<br>p=0.7167 | | F (1, 57) = 0.5149<br>p=0.4780 |
| fast $\Delta V_m$ vs baseline | F (1, 57) = 2.576<br>p=0.1140 | **F (1, 57) = 4.462**<br>**p=0.0390** | | F (1, 57) = 0.002304<br>p=0.9619 |
| fast $\Delta V_m$ latency | **F (1, 57) = 6.870**<br>**p=0.0112** | F (1, 57) = 0.3350<br>p=0.5650 | | F (1, 57) = 1.683<br>p=0.1998 |
| slow $\Delta V_m$ vs baseline | F (1, 57) = 1.203<br>p=0.2774 | F (1, 57) = 2.232<br>p=0.1407 | | F (1, 57) = 1.291<br>p=0.2606 |
| slow $\Delta V_m$ latency | F (1, 57) = 0.02167<br>p=0.8835 | F (1, 57) = 1.193<br>p=0.2793 | | F (1, 57) = 0.9738<br>p=0.3279 |
| Fisher's LSD analysis of significant two-way ANOVA |  |  |  |  |
|  | 3wk VEH<br>vs.<br>3wk PNA | adult VEH<br>vs.<br>adult PNA | 3wk VEH<br>vs.<br>adult VEH | 3wk PNA<br>vs.<br>adult PNA |
| fast $\Delta V_m$ vs baseline | p=0.1400 | p=0.1414 | p=0.2564 | p=0.2650 |
| fast $\Delta V_m$ latency | p=0.1990 | p=0.6058 | p=0.3338 | **p=0.0099** |

The 6-min apamin treatment changed the post-spike-train membrane potential trajectory in all groups, unmasking a fast depolarization that peaked within 300ms of the end of current injection (Figure 8A, red trace). GnRH neurons from 3wk-old PNA mice had the largest post-spike-train depolarization amplitude; this was decreased in adulthood to values observed in VEH mice (Figure 8B, two-way ANOVA age p=0.0226, treatment x age p=0.0280, Fisher’s LSD 3wk VEH vs 3wk PNA p<0.05, 3wk PNA vs adult PNA p<0.01, Table 14). PNA treatment shortened the latency to peak of the depolarization (Figure 8C, two-way ANOVA, PNA treatment p=0.017), with the difference being more pronounced between adult PNA and adult VEH group (Fisher’s LSD, p<0.05). To minimize the influence of noise and prolonged tails when quantifying the AUC and duration of the post-spike-train depolarization, the onset and termination were defined as the point when potential crossed 10% of the depolarization peak value (AUC at 10% maximum, duration at 10% maximum). There was no difference in post-spike-train depolarization AUC or duration at 10% maximum among groups.

**Figure 8.**
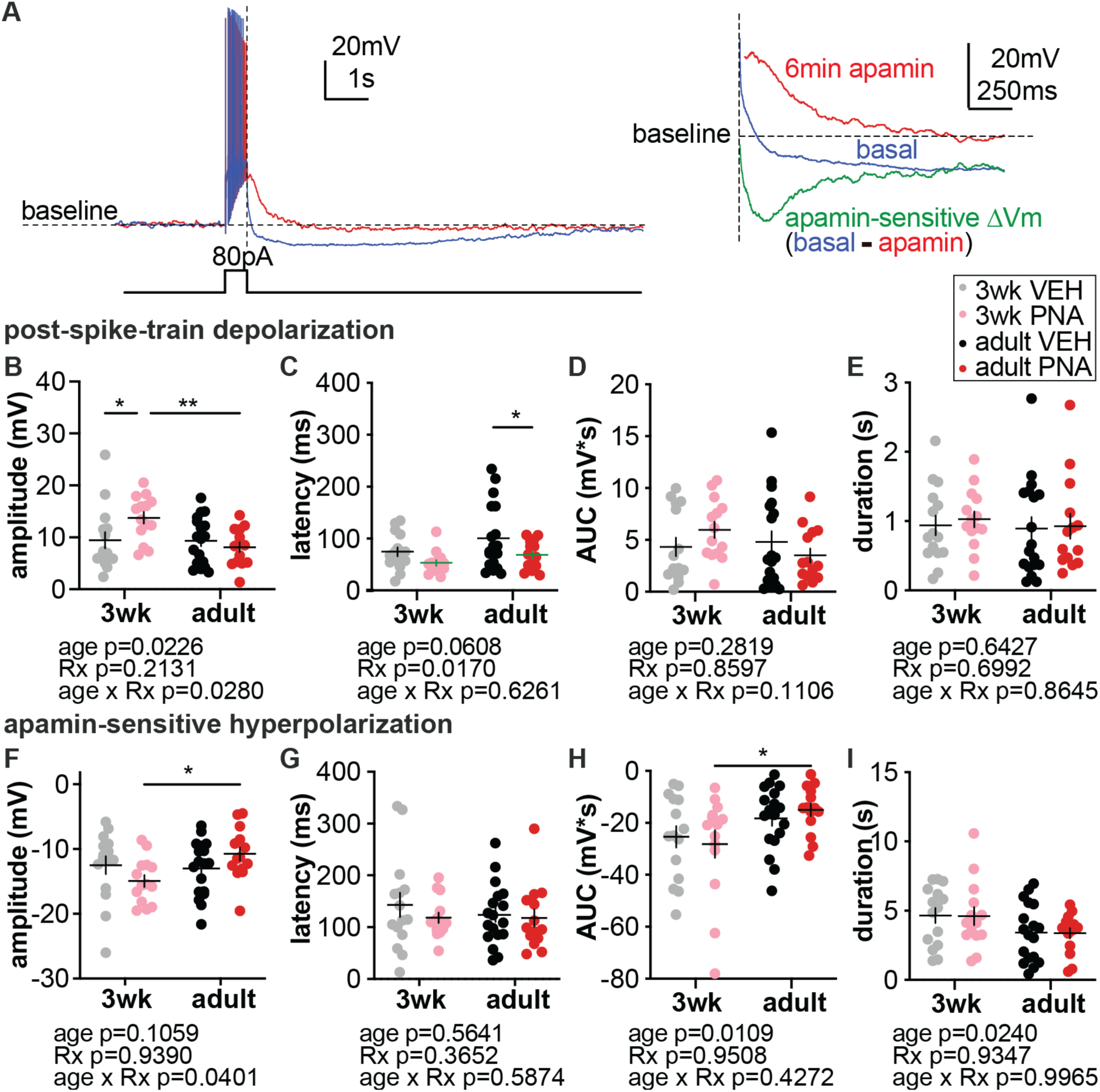
The post-spike-train depolarization unmasked by apamin, and apamin-sensitive SK-mediated hyperpolarization. **A**, left: representative membrane potentials from the same GnRH neuron in response to 80pA current injection under basal conditions (blue) and after 6min of apamin treatment (red). Right: enlargement of the figure on the left showing post-spike-train membrane potentials under basal conditions (blue), after apamin treatment (red) and the SK-mediated hyperpolarization (green) obtained by subtracting the red trace from the blue trace. Horizontal dashed line: baseline potential; vertical dashed line: end of current injection. **B-E**, post-spike-train depolarization peak value (B), latency (C), AUC at 10% maximum (D), and duration at 10% maximum (E). F-I, SK-mediated hyperpolarization peak amplitude (F), latency (G), AUC at 10% maximum (H), and duration at 10% maximum (I). RX, prenatal treatment. *p<0.05, **p<0.01, Fisher’s LSD.

**Table 14.** Two-way ANOVA of the effects of apamin on post-spike train change in membrane potential (Figure 8).

**Post-spike-train depolarization**
| property | age |  | treatment |  | interaction |
| --- | --- | --- | --- | --- | --- |
| amplitude | F (1, 57) = 5.495<br>p=0.0226 |  | F (1, 57) = 1.586<br>p=0.2131 |  | F (1, 57) = 5.087<br>p=0.0280 |
| latency | F (1, 57) = 3.658<br>p=0.0608 |  | F (1, 57) = 6.045<br>p=0.0170 |  | F (1, 57) = 0.2400<br>p=0.6261 |
| AUC | F (1, 57) = 1.180<br>P=0.2819 |  | F (1, 57) = 0.03154<br>p=0.8597 |  | F (1, 57) = 2.627<br>p=0.1106 |
| duration | F (1, 57) = 0.2176<br>p=0.6427 |  | F (1, 57) = 0.1508<br>p=0.6992 |  | F (1, 57) = 0.02939<br>p=0.8645 |
| Fisher's LSD analysis of significant two-way ANOVA |  |  |  |  |  |
|  | 3wk VEH<br>vs.<br>3wk PNA | adult VEH<br>vs.<br>adult PNA | 3wk VEH<br>vs.<br>adult VEH | 3wk PNA<br>vs.<br>adult PNA |  |
| amplitude | p=0.0181 | p=0.4747 | p=0.9482 | p=0.0027 |  |
| latency | p=0.1780 | p=0.0375 | p=0.0822 | p=0.3367 |  |

**Apamin-sensitive hyperpolarization**
| property | age | treatment |  | interaction |
| --- | --- | --- | --- | --- |
| amplitude | F (1, 57) = 2.700<br>p=0.1059 | F (1, 57) = 0.005912<br>p=0.9390 |  | **F (1, 57) = 4.413**<br>**p=0.0401** |
| latency | F (1, 57) = 0.3365<br>p=0.5641 | F (1, 57) = 0.8331<br>p=0.3652 |  | F (1, 57) = 0.2977<br>p=0.5874 |
| AUC | **F (1, 57) = 6.929**<br>**p=0.0109** | F (1, 57) = 0.003836<br>p=0.9508 |  | F (1, 57) = 0.6396<br>p=0.4272 |
| duration | **F (1, 57) = 5.379**<br>**p=0.0240** | F (1, 57) = 0.006762<br>p=0.9347 |  | F (1, 57) = 1.945e-005<br>p=0.9965 |
| Fisher's LSD analysis of significant two-way ANOVA |  |  |  |  |
|  | 3wk VEH<br>vs.<br>3wk PNA | adult VEH<br>vs.<br>adult PNA | 3wk VEH<br>vs.<br>adult VEH | 3wk PNA<br>vs.<br>adult PNA |
| amplitude | p=0.1369 | p=0.1492 | p=0.7373 | **p=0.0135** |
| AUC | p=0.6113 | p=0.5361 | p=0.1826 | **p=0.0229** |
| duration | p=0.9572 | p=0.9503 | p=0.0937 | p=0.1190 |

The change in post-spike-train membrane potential following apamin perfusion indicates that the apamin-sensitive SK current shapes the membrane potential after a train of action potentials, counteracting depolarization and facilitating repolarization/hyperpolarization. To assess the net effect of apamin-sensitive SK current on post-spike-train membrane potential, we subtracted the membrane potential trace recorded 6min with apamin perfusion from that recorded in the basal state before treatment. The subtraction revealed a SK-mediated hyperpolarization that peaked within 350ms of the end of current injection (Figure 8A, green trace). GnRH neurons from 3wk-old PNA mice exhibited the largest SK-mediated hyperpolarization; this declined in adulthood (Figure 8F, two-way ANOVA age x treatment p=0.0401; Fisher’s LSD 3wk PNA vs adult PNA p<0.05). Similarly, AUC of the SK-mediated hyperpolarization at 10% maximum in GnRH neurons from PNA mice declined in adulthood (Figure 8H, two-way ANOVA, age p=0.0109; Fisher’s LSD 3wk PNA vs adult PNA p<0.05). There was no difference in the latency to the SK-mediated hyperpolarization peak (Figure 8G). Compared with 3wk groups, GnRH neurons from both adult VEH and PNA mice had shorter duration of SK-mediated hyperpolarizations at 10% maximum (Figure 8I, two-way ANOVA age p=0.024).

## Discussion

Precise control of the frequency of GnRH pulses is required to drive the female reproductive cycle. In women with hyperandrogenemic PCOS, typical frequency changes in LH pulses, which reflects GnRH pulses, are replaced by a persistent high frequency pulses. Similarly, LH-pulse frequency is elevated in PNA mice, and these mice lack the developmental changes in GnRH neuron firing activity observed in VEH mice. Here we examined age and PNA-induced changes in voltage-gated Ca^2+^ currents, which are critical to excitation-secretion coupling (Ludwig et al., 2002; Chen and Moenter, 2023), and firing dynamics (Lee et al., 2010; Weiss and Zamponi, 2024; Esquivel-Garcia et al., 2025). Calcium currents were altered in PNA mice and from prepubertal to adult animals, indicating they may contribute to the phenotypes in PNA mice and possibly women with PCOS.

The present results support prior studies of GnRH neuron I_Ca_ that report the total current is comprised of medium-slow and fast inactivating components. The medium/slow I_Ca_ displays slow voltage-dependent inactivation characteristic of L-type Ca^2+^ channels (Stotz and Zamponi, 2001; Striessnig et al., 2004; Catterall, 2011), whereas the fast I_Ca_ component is likely mediated by N, P/Q, and R Ca^2+^ channels, with a minimal T-type component. Our data extend prior studies by demonstrating that GnRH neurons from PNA mice exhibited increased I_Ca_ density compared with VEH groups at both prepubertal and adult ages for total I_Ca_, as well as both the medium-slow and fast subcomponents. The among-group differences were evident at depolarized membrane potentials consistent with activation of high-voltage-activated calcium channels. In contrast, no differences were observed at more hyperpolarized potentials that would tend to activate low-voltage-activated T-type currents. Minimal current activation at those potentials is consistent with prior reports (Kato et al., 2003; Nunemaker et al., 2003; Sun et al., 2010) and with reports that Ca_V_3 subunits encoding T-type channels are expressed only in a subset of GnRH neurons (Zhang et al., 2009). Altered T-type I_Ca_ is thus not likely a major cause of development or PNA-induced changes in GnRH neuron activity (Dulka and Moenter, 2017).

I_Ca_ properties depended on both PNA treatment and age. GnRH neurons from VEH mice exhibited a developmental depolarization in voltage-dependent activation of total Ca^2+^, likely attributable to the medium-slow component, that was absent in PNA mice. The more hyperpolarized activation of Ca^2+^ channels in GnRH neurons from 3wk-old VEH mice may help sustain membrane depolarization near action potential threshold (Striessnig et al., 2014) and could thus in part explain increased spontaneous action potential firing in this group relative to 3wk PNA or adult groups (Dulka and Moenter, 2017). PNA mice lack developmental depolarization of Ca^2+^ current activation and exhibited no changes in spontaneous firing activity with age. In contrast to shifts in activation, which were exclusive to VEH mice, time-dependent inactivation of the fast I_Ca_ was slower in adult mice regardless of PNA treatment. This would allow more Ca^2+^ entry in GnRH neurons from adult mice, favoring neuropeptide release (Catterall, 2011) and possibly contributing to the pubertal transition. GnRH neurons from PNA mice had a depolarized V_0.5inact_ compared with VEH groups at both ages, which could increase Ca^2+^ entry in GnRH neurons to facilitate neuropeptide release. Alterations in expression or posttranslational modifications of Ca channel α and/or Δ subunits could influence these properties.

Steroid milieu changes with both development and PNA and could underlie some of the observed shifts in I_Ca_ properties. More L-type but less N-type mediated currents were reported in 4–10-day old mice compared with adults (Nunemaker et al., 2003). *In vivo* estradiol treatment of adult ovariectomized mice at a level sufficient to induce a preovulatory GnRH/LH surge increases the expression of Ca_V_1.3, Ca_V_2.2, and Ca_V_2.3 channels in GnRH neurons, but had no effect on Ca_V_1.2 (Bosch et al., 2013). Ovariectomized female mice treated with an estradiol leading to diurnal changes in GnRH neuron firing rate also exhibit diurnal changes in HVA I_ca_ amplitude with alterations in L-and N-type subtypes (Sun et al., 2010). Removing steroid negative feedback by castration increases Ca^2+^ currents in GnRH neurons from both male (Chen and Moenter, 2023) and female mice (Sun et al., 2010). Perhaps most relevant, treatment of ovariectomized mice with DHT increased I_Ca_ (Sun and Moenter, 2010) and PNA mice have elevated testosterone levels (Sullivan and Moenter, 2004; Moore et al., 2013). Shifts in steroid milieu, particularly androgens, could thus mediate observed changes in I_Ca_.

GnRH neuron excitability was examined to assess functional changes in GnRH neuron output that the observed changes in I_Ca_ may contribute to, using an EGTA-free pipette solution to minimize buffering. Firing rate in cells from 3wk VEH mice was greater than adults; this developmental change was not evident in cells from PNA mice. These excitability changes resemble age and PNA-related changes in extracellularly-recorded spontaneous firing activity of GnRH neurons (Dulka and Moenter, 2017). In studies using EGTA-containing pipette solution, GnRH neurons from VEH and PNA mice had comparable excitability that increased with age (Jaime and Moenter, 2022). Differences with and without buffering suggested I_Ca_ and [Ca^2+^]_in_ play roles in GnRH neuron excitability in VEH and PNA mice during development. In the present work, action potential threshold was hyperpolarized in 3wk vs adult mice, largely attributable to changes in 3wk VEH mice, consistent with the firing changes observed. Age also altered the afterhyperpolarization (AHP), with increased amplitude and a later peak time observed in adults. Since similar AHP changes were observed in studies using buffered and unbuffered pipette solution, Na^+^ (threshold) and K^+^ currents (AHP) likely mediate these changes in first action potential properties.

During an action potential burst, increased [Ca^2+^]_in_ can open Ca^2+^-activated K^+^ channels, a mechanism that could regulate GnRH neuron excitability by increasing intraspike intervals and thereby reducing firing rate. Blockade of SK channels with apamin increased GnRH neuron firing rate in all groups except PNA adults. Apamin also affected the membrane hyperpolarization after termination of the current injections, confirming a prior report of a role for a rapidly activating apamin-sensitive SK current (Lee et al., 2010). In our recordings under basal conditions, the post-spike-train fast ΔV_m_, which peaks within 150ms after the termination of current injection, was shifted towards zero or became depolarizing in GnRH neurons from PNA mice at both ages. This reduction in hyperpolarization may provide a transient window favoring another spike or change the availability of voltage-dependent channels.

Blocking SK channels with apamin unmasked a post-spike-train depolarization, indicating the ΔV_m_ arises from a competition between depolarizing and hyperpolarizing currents, including SK. Afterdepolarizations are mediated by a TTX-sensitive sodium current in GnRH neurons (Chu and Moenter, 2006) or calcium current in magnocellular neuroendocrine neurons (Andrew and Dudek, 1983, 1984; Li and Hatton, 1996). In GnRH neurons, the afterdepolarization favors increased spontaneous firing. Interestingly, GnRH neurons from 3wk PNA mice exhibited the largest post-spike-train depolarization that is counterbalanced by a stronger the SK-mediated hyperpolarization. The stronger SK-mediated hyperpolarization constrains firing and is consistent with their increased firing rate in response to apamin. It may also help explain the lower firing rate of GnRH neurons from 3wk PNA versus 3wk VEH mice observed in extracellular and current-clamp recordings. The stronger SK-mediated hyperpolarization may be an intrinsic homeostatic compensatory mechanism to counteract the increased excitatory GABAergic drive these cells receive at this age (Berg et al., 2018). In adulthood, the post-spike-train depolarization in GnRH neuron from PNA mice decreased to levels comparable to VEH mice, and this is accompanied by decreased SK-mediated hyperpolarization. In contrast, GnRH neurons from VEH mice had stable SK-mediated hyperpolarization during development. It should be noted that the diminished SK-mediated hyperpolarization in adult PNA GnRH neurons appeared despite increased I_Ca_. suggesting reduced coupling of increased I_Ca_ to opening of SK channels. The reduced SK-mediated hyperpolarization in adult PNA GnRH neurons is consistent with their blunted firing rate response to apamin. GnRH neurons from adult PNA mice thus have reduced capacity to constrain firing with this mechanism, and any compensation it provided for increased excitatory GABAergic drive, which persists into adulthood in PNA mice, is lost.

Together these findings demonstrate that I_Ca_ in GnRH neurons is modified both by developmental stage and PNA treatment. These changes may contribute to the normal pubertal transition, as well as transitory compensation mechanisms involving calcium-activated potassium currents for increased excitatory drive and their ultimate failure in PNA mice. Future studies of the role of I_Ca_ in GnRH neuron firing dynamics can be approached with dynamic clamp, whereas its role in excitation secretion coupling can be examined with real-time measures of GnRH release with fast-scan cyclic voltammetry.

## Conflict of interest

The authors declare no competing financial interests.

## Acknowledgements

We thank Elizabeth Wagenmaker, Laura Burger for expert technical support and James L. Kenyon, University of Nevada, Reno, for the Excel spreadsheet used to calculate junction potentials. Supported by National Institute of Health/*Eunice Kennedy Shriver* National Institute of Child Health and Human Development R01HD104345.

## Abbreviations

GnRH: gonadotropin-releasing hormone;
LH: luteinizing hormone; FSH, follicle-stimulating hormone;
GFP: green fluorescent protein;
VEH: prenatal vehicle treatment;
PNA: prenatal androgen treatment;
DHT: dihydrotestosterone

## Notes

### Competing Interest Statement

The authors have declared no competing interest.

